# Hepatocellular carcinoma hosts immature neurons and cholinergic tumors that correlate with adverse molecular features and outcomes

**DOI:** 10.1101/2022.06.14.495889

**Authors:** Charlotte A. Hernandez, Claire Verzeroli, Ievgeniia Chicherova, Abud-José Farca-Luna, Laurie Tonon, Pascale Bellaud, Bruno Turlin, Alain Fautrel, Zuzana Macek-Jilkova, Thomas Decaens, Sandra Rebouissou, Alain Viari, Fabien Zoulim, Romain Parent

**Affiliations:** Pathogenesis of Chronic Hepatitis B and C laboratory - LabEx DEVweCAN, Inserm U1052, Cancer Research Centre of Lyon, F-69003 Lyon, France, University of Lyon, F-69003 Lyon, University of Lyon 1, ISPB, Lyon, F-69622, France, CNRS UMR5286, F-69083 Lyon, France, Centre Léon Bérard, F-69008 Lyon, France; Fondation Synergie Lyon Cancer, Plateforme de bioinformatique Gilles Thomas, Centre Léon Bérard, F-69008 Lyon, France; H2P2 platform, University of Rennes, Rennes, France; Institute for Advanced Biosciences, Inserm U1209, University of Grenoble-Alpes, F-38700 La Tronche, France; Service d’hépato-Gastroentérologie, Pôle Digidune, CHU Grenoble-Alpes, 38700 La Tronche, France; Centre de Recherche des Cordeliers, Inserm, Sorbonne Université, USPC, Université Paris Descartes, Université Paris Diderot, Paris, France; Hospices Civils de Lyon, Service of Hepato-Gastroenterology, F-69001 Lyon, France

**Author notes:** Equal contributions (Charlotte A. Hernandez, Claire Verzeroli). **Authors contributions:** CAH, CV, IC, AJFL, PB, ZMJ and RP generated and analyzed experimental data. PB, AF and ZMJ developed critical approaches and methods for the study. LT, BT, AF, ZMJ, TD, SR, AV, FZ and RP critically amended and conceptually enriched the paper. RP wrote the paper.

**Keywords:** HCC, autonomic nervous system, IHC, transcriptomics, cholinergic

## Abstract

**Background & aims:** The unexplained interpatient variation in hepatocellular carcinoma (HCC) remains a major challenge. We aimed at addressing the under-explored association between the disease and the hepatic autonomic nervous system (ANS).

**Methods & Results:** We in-depth characterized the innervation of French biobanks HCC samples by conventional biochemistry methods. We also applied bioinformatics approaches to the TCGA dataset in order to stratify samples according to neural features and molecular correlates. We highlighted the predominant parasympathetic polarity of HCC nerves, and demonstrated that a cirrhotic rat model of aggressive HCC hosts liver neurogenesis with cholinergic features. Using the TCGA dataset, we then defined an HCC neural signature, derived from adrenergic and cholinergic receptor levels, that allowed patient stratification into two classes. Cholinergic tumors correlated with *TP53* mutations (*p* ≤ 0.05), shorter progression-free interval (PFI) and overall survival (OS), displayed more pathogenic molecular traits (*e.g*., AFP-rich, proliferative tumors, mitotic functions including DNA repair, EMT, Ras, and Akt/mTOR pathways), aggressive HCC signatures and B cell accumulation. Instead, adrenergic tumors, predominant in patients aged >60 and with mutated *CTNNB1*, were correlated with better OS and PFI (*p* < 0.05), and numerous immune pathways.

**Conclusions:** Our results depict neural features of HCC and how the existing tumor classification may also be shaped by neural inputs. Altogether, we show that the parasympathetic branch of the ANS is implicated in the pathobiology of HCC, and advocate for the use of ANS-targeting drugs in HCC research, many of which are clinically safe and well characterized.

## Introduction

Primary liver cancer casualties are ranked 3^rd^ worldwide (1) and are still on the rise despite the recent advent of adequate hepatitis B and C (HBV and HCV) therapies. Genetic diseases of the liver and hepatic comorbidities, such as alcoholic liver disease (ALD) and metabolic syndrome with non-alcoholic steato-hepatitis (NASH), are long-term cooperators or independent factors fostering the onset of HCC and enhancing disease heterogeneity. Though HCC is known to develop in 90% of cases of cirrhosis (2), its onset and clinical outcomes, in terms of phenotypes and speed of progression, are highly variable from one patient to another. Despite the identification of several potential therapeutic targets, most drugs have failed to exceed the efficacy of currently available compounds. Treatments with tyrosine kinase inhibitors (TKIs) for instance lead to short-term, unavoidable relapse (3), whereas treatment with immune check-points inhibitors or growth factors inhibitors currently provide some hope for only a minority of patients with unresectable HCC (2).

In this respect, cellular/tissular structures linking the general pathophysiology of the patient with HCC may be of interest, as they are patient-specific and may uncover novel ways of defining stratification criteria. In line with such notions, several recent original papers ^(4),(5),(6),(7, 8)^ and related commentaries (9, 10) highlighted the relevance of studying cancer neurosciences of peripheral organs. In that context, pathological innervation and ANS involvement or dysregulation have been identified in ovarian (4), prostate (5), gastric (6) and pancreatic (7, 8) cancers, nurturing tumor stroma and conferring stronger carcinogenic properties. Moreover, ANS post-synaptic receptors have been shown to be favorably actionable in some experimental conditions in cancer (5, 7, 8).

The autonomic nervous system (ANS) comprises the sympathetic (adrenergic signaling) and parasympathetic (cholinergic signaling) arms that relay signals both ways along the brain-liver neural axis in order to regulate involuntary functions of the body by adjusting its internal functions, after an external stimulus. The liver is an innervated organ that hosts autonomic afferent and efferent ANS nerves, in constant communication with the central nervous system (CNS) through the brainstem (11). Afferent and efferent nerves are made of adrenergic (relies on epinephrine or norepinephrine as its neurotransmitter, stress signal) and cholinergic (relies on acetylcholine as its neurotransmitter, resting signal) fibers that each convey mediators to regulate liver functions in real-time.

As a consequence, as also notably pointed out by Tracey’s theory and evidence for cholinergic blockade of inflammation {Pavlov, 2015 #7355}, these signals also regulate several processes that may directly or indirectly impact HCC onset and growth. However, data on the association between HCC and neural factors are scarce and sometimes conflicting. It was reported that portal hypertension, a recognized risk factor for HCC development and recurrence (12, 13), is correlated with ANS dysfunction (14). In addition, proliferation of hepatocytic progenitors, instrumental in HCC, is impaired by adrenergic signaling (15). Conversely, cholinergic signaling was shown to attenuate apoptosis in the mouse liver (16, 17), and liver angiogenesis is under positive sympathetic regulation (18). Interestingly, human liver ANS innervation is more developed than in rodents. Indeed, it extends deeper into the lobule (11), increasing its capacities of regulation. This latter notion suggests that ANS-related mechanisms observed in animals may play more important roles in humans.

Here, we characterized several ANS markers of HCC samples by Western blot (WB) and immunohistochemistry (IHC), unveiling a predominant parasympathetic signaling compared to normal liver tissue. Considered by bioinformatics, the definition of two neural HCC classes based on levels of adrenergic or cholinergic receptor transcripts, one cholinergic and one adrenergic, enabled the stratification of tumors with more or less aggressiveness, respectively. Our findings shed light on, as yet, unexplored characteristics of the disease and thus offer a realistic outlook for considering alternative HCC investigations and treatment strategies, given the diversity and safety of currently approved ANS-targeting drugs.

## Materials and Methods

Sections related to rat liver samples, western blotting, total RNA processing, immunohistochemistry and statistics are provided in **Supplementary Information 1**.

### Clinical liver samples

Hepatocellular carcinoma samples used in this study were from the French Liver Biobanks network (INCa, BB-0033-00085, under IRB agreement of Inserm Ethics Committee (CEEI, #12-063) and the TCGA Research Network (https://www.cancer.gov/tcga) HCC-LIHC cohort (19). The table presenting data expression was downloaded with the tool TCGA biolinks http://bioconductor.org/packages/release/bioc/html/TCGAbiolinks.html Data were crossed with previously reported metadata to obtain a cohort of 193 patients. Normal liver samples (French South-East region IRB agreement #A16-207) obtained from safety margins of hepatic resections of colorectal cancer metastasis, were histologically normal and devoid of HBV or HCV infection.

### Neuronal receptor score calculation and cohort classification

ssGSEA scores were computed through clusterProfiler R package. Gene set scores were calculated with the ssgsea method (20) from the GSVA package (21). ssGSEA is a variation of Gene Set Enrichment Analysis (GSEA) that calculates enrichment scores for each pairing of gene set and sample. ssGSEA transforms data to a higher space level, from individual genes to genes sets (a list of functionally related genes) or pathways. As other methods, this analysis identifies enriched or overrepresented gene sets among a list of ranked genes providing the advantage that ssGSEA enrichment scores reflect the activity level of the biological process in which the genes of a particular gene set are coordinately up- or down-regulated within a sample. This transformation allows the interpretation of cell status through activity levels of biological processes and pathways rather than through the expression levels of individual genes. Here, two ssGSEA scores were calculated from both signatures, one including all adrenergic receptor genes, and the other including all cholinergic receptor genes in order to obtain an adrenergic score (AEs) and a cholinergic score (ChEs), respectively. To obtain a global neurotransmitter receptor score (NRs), the difference based on the adrenergic score minus the cholinergic score was calculated for each sample. Samples were split in two classes defined by receptor expression: those with higher NRs than median were named adrenergic while those with lower NRs than median were named cholinergic.

### Differential Gene Expression

Differential gene expression was performed with DESeq2 Bioconductor R package (22) using as contrast previous clustering (classes by synaptic receptor, adrenergic-cholinergic signature). padj was set to 0.01 and Log2 Fold Change abs > 0.58. “High” (adrenergic) class was considered as reference.

### Pathway enrichment analysis

All pathway-enrichment analyses were conducted using MSigDB gene sets H, C2, and C5 from msigdbr R package (23). Overrepresentation analysis of pathways was performed with enricher from clusterProfiler (24) using as input over-expressed and down-expressed genes previously obtained by differential gene expression analysis.

### Survival analysis

The survival data (OS, PFI) were extracted from Liu et al. (25). Survival analyses were conducted with survival R packages (26, 27) and survminer (28). The survival analysis was performed at 2, 3, 5 and 10 years. p < 0.05 was considered significant.

### Microenvironmental analysis

Estimation of immune and non-immune cell fractions from the tumor microenvironment were determined through gene expression analysis using Immunedeconv R package (29) together with xCell enrichment method (30). This method allows sample to sample comparisons and to determine stroma, immune and microenvironment scores.

## Results

### Neurogenesis of parasympathetic orientation in a cirrhosis-associated HCC rat model

Emerging evidence suggests a potential dysregulation of ANS signaling in several cancers in humans and in animal models (4–8). To our knowledge, neuroregulation in HCC has never been investigated *in vivo*. HCC occurs in a cirrhotic background in 90% of cases. A rat model that allows HCC growth on genuine cirrhosis, a near obligatory step for HCC onset in the clinic, has been extensively characterized (31) (also in **Suppl. Figure 1** herein) and shows documented clinical relevance in particular with respect to the proliferative class of HCC (32). In this context, we herein evaluated the ability of this diethylnitrosamine (DEN)- treated HCC rat model to recapitulate such processes (4–8). The methodology used for the experimental induction of HCC is shown in **Figure 1A.** To characterize HCC innervation, the following classically validated neuron markers were considered: NeuN (phospho- and total RBFOX3) as a mature neuron marker, DCX (phospho- and total forms) and INA as immature neurons-specific markers, TH (TYR3H) and VAChT (SLC18A3) as specific ANS neuron markers for sympathetic and parasympathetic neurons, respectively. Total NeuN signals increased throughout disease progression. Interestingly, signals related to the DCX progenitor marker increased transiently, yet sharply, in samples harboring cirrhosis and HCC small nodules (shown in **Suppl. Fig. 1**). In the case of ANS-specific markers, though levels of the TH marker (adrenergic) remained unaltered throughout disease progression, an increase in the cholinergic VAChT marker was observed in rats suffering from HCC. Degradation of *β*-tubulin was correlated with DEN-treatment and was likely derived from hepatic cytolysis and release of cytosolic contents, due to DEN-related, neoantigen-targeting, immune activation (**Figure 1B**). Intriguingly, along this line, a tight correlation was observed between DCX expression and *β*-tubulin degradation throughout progression of HCC-predisposing chronic liver disease (CLD) (**Figure 1C**). This suggests that HCC neural remodeling occurs as a consequence of cytolysis or parenchymal remodeling, as recently demonstrated in NASH (33). This prompted us to analyze the quantitative evolution of neural markers during CLD and HCC progression, which confirmed neo-neurogenesis, and the parasympathetic neural features of HCC in the rat (**Figure 1D-H**), features pertinent to the human phenotype described herein. Of note, consistently, this model was recently shown to recapitulate features of clinical lesions of poorer outcomes (32). Altogether, such data indicate that alteration hepatic neural features correlate with progression towards HCC *in vivo*.

**Figure 1.**
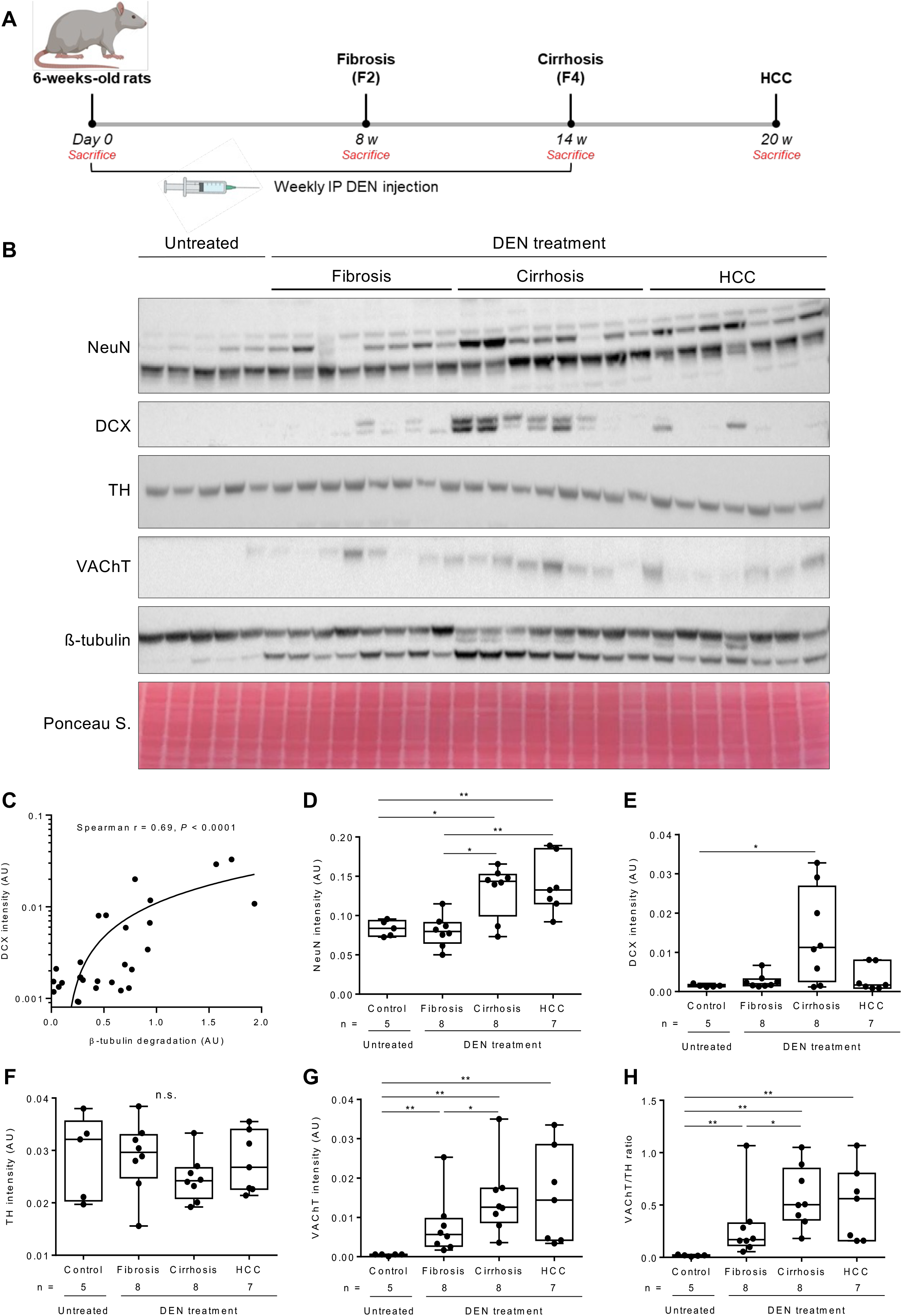
Evolution towards HCC and hepatic remodeling are correlated with neurogenesis and parasympathetic orientation in the cirrhotic rat. **A.** Experimental outline. 6-week-old Fischer 334 male rats were given DEN at 50 mg/kg weekly from day 0 to week 14, allowing progression of chronic liver disease to fibrosis, cirrhosis, decompensated cirrhosis and HCC. **B.** Immunoblotting of total NeuN (mature neurons), total DCX (immature neurons), TH (adrenergic) and VAChT (cholinergic) neural markers. **C.** DCX induction is correlated with parenchymal remodeling. DCX levels were plotted against ratios of full-length versus degraded tubulin signals shown in panel B. Spearman test (*** p < 0.001). **C-H.** Signal quantification was done using the Fiji software on non-saturated images (as detailed in the *Materials and Methods* section, also in all subsequent data), prior to statistical plotting using the Mann-Whitney test (* p < 0.05, ** p < 0.01). Five to eight rats were used per time point.

### Infiltration of neural progenitors of parasympathetic orientation in human HCC samples

Given that the human liver ANS innervation extends deeper into the lobule (11) than in rodents, the extent of involvement of neural processes in human HCC may be substantial. We then investigated by immunoblotting the presence of ANS neural markers in paired peritumoral (minimum distance of 2 cm) and tumoral clinical HCC samples. These samples, obtained from the French National HCC biobank, were evenly distributed across the main four HCC etiologies (HBV, HCV, ALD, NASH; 24-26% each). The main characteristics of the patients are provided in the **Suppl. Table 1**. For optimal comparability between samples of each etiology, the HBV RNA and HCV RNA intrahepatic viral loads were limited to a maximum of 2 log_10_ difference.

We first considered by immunoblotting the presence of ANS neural markers in normal livers (both uninfected and F0, see Materials & Methods) versus HCC samples. Western blotting highlighted specific DCX and INA positivity in HCC samples, with a slight decrease in mature neural marker NeuN expression in HCC, indicating the presence of immature neurons. In addition, HCC samples displayed a profound depletion of the adrenergic marker TH at the benefit of the cholinergic neural marker VAChT (**Figure 2A**), prompting further analysis. We then compared the expression levels of such markers after thorough technical validation of the antibodies of interest (**Suppl. Fig. 2A-B**) at the peritumoral (cirrhotic/F4 stage) and tumoral levels in samples of the main HCC etiologies ((HBV (n = 14), HCV (n = 9), ALD (n = 14) and NASH (n = 14), total of 51 patients). The intensity of neuron markers was heterogeneous between etiologies. Only two clinico-biological features were significantly different between peritumoral and tumoral tissue: DCX enrichment in HBV+ samples and TH depletion in NASH samples (p < 0.05). When focusing on tumor differentiation, we then observed that well differentiated HCC samples were inversely correlated with mature innervation (NeuN+) in HBV-related tumors (**Suppl. Fig. 2C-E**). No other marked difference could be revealed between peritumoral and tumoral samples (**Suppl. Fig. 3-7**). Overall, the important heterogeneity in neural markers observed across all tumors (significance is summarized in **Suppl. Table 2**), reflects, and maybe even affects, the well-known heterogeneity of HCC. Our data indicate that neo-neurogenesis occurs in all HCC etiologies, and that NASH is the primary etiology able to drive associations with neural markers. Importantly, such data also indicate that neurogenesis and its related alterations precedes HCC *per se* since no significant difference in the expression of markers could be observed in several instances between F4 and HCC.

**Figure 2.**
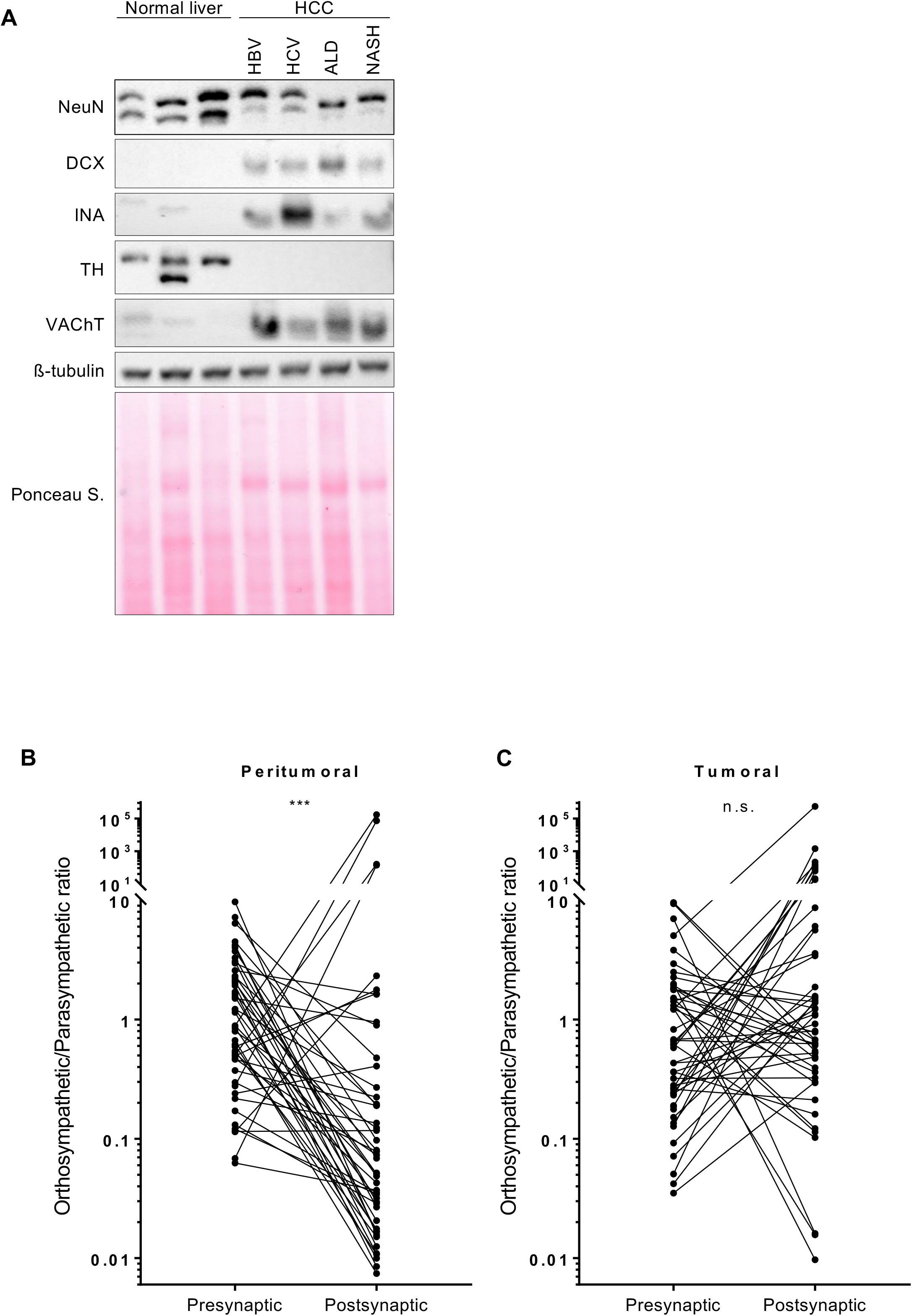
Expression of mature and progenitor neural markers in human HCC. **A.** Immunoblotting of NeuN (mature neurons), DCX and INA (immature neurons), TH (adrenergic) and VAChT (cholinergic) neural markers carried out on normal liver and HCC samples. **B-C.** Tumor-specific uncoupling between neuron marker and post-synaptic receptors upon comparison with non-tumoral paired tissues. Relationship between neuronal pre-synaptic marker and post-synaptic marker expression. TH and VAChT expression levels in peri-tumoral and tumoral samples (n = 51 HCC cases) were evaluated. Gene expression levels of adrenergic and cholinergic receptors was evaluated. Orthosympathetic/Parasympathetic ratios were calculated for each patient taking into consideration TH over VAChT signals and (*ADRA1A+ADRB2+ADRA1B*) over (*CHRNA4+CHRNA7+CHRM3*) signals, where those six targets correspond to the most expressed of both categories in patient samples. Mann-Whitney test (* p < 0.05, ** p < 0.01, *** p < 0.001).

Next, we sought to gain insight into the localization of neural signals in human samples. A set of 24 HCC patient samples derived from the four main etiologies ((HBV (n = 7), HCV (n = 4), ALD (n = 9), NASH (n = 4)) were subjected to Masson’s trichrome staining to expose tissue architecture, and to NeuN, DCX, TH and VAChT immunostaining coupled with DAPI staining. INA signals could not be validated by IHC. The technical validations for NeuN, DCX, TH and VAChT staining are provided in the **Suppl. Fig. 8.** As a first attempt to document localization of these antigens, we performed standard 2D staining of a pilot cohort of 24 tumors. TH staining was negligible in both frequency and intensity throughout samples (**Suppl. Fig. 9A-F**). Importantly, neuronal markers DCX and VAChT could be found in the tumor bulk where they co-stained (**Suppl. Fig. 9G-J**), NeuN was never detected in the tumor bulk, suggesting that intra-tumoral neurogenesis is more immature and dynamic than its capsular counterpart. As observed by WB, the predominant ANS co-labeling was specific for immature DCX+ fibers and parasympathetic neurons, suggesting that HCC neo-neurogenesis is largely parasympathetic, a notion in line with previous data on NASH (33). Such observations are to be confirmed with for instance 3D staining after tissue clarification for better morphological insight. They were unrelated to any specific HCC etiology, and however in line with findings on other solid malignancies (4–8) that these tumors host nerves with migrating potential, likely enabling their interaction with post-synaptic receptors.

### Modulation of ANS receptor transcripts from cirrhosis to HCC

Transduction of neural signals in the diseased tissue implicate postsynaptic ANS receptors. As a consequence, we compared the expression of several transcripts encoding ANS receptors in paired cirrhosis/F4 and HCC samples in the same cohort (n = 161 patients) from the French National HCC biobank. Neuronal (presynaptic) mRNA markers such as *RBFOX3, DCX, TYR3H*, and *SLC18A3* could not be amplified by RT-qPCR using ad hoc neuronal control cells (not shown), likely due to limited amounts of mRNAs in the distal, hepatic extremity of the axon, prompting for the consideration of TH over VAChT ratios at the presynaptic level. Post-synaptic neural markers belonging to detectable and well-characterized factors from the adrenergic group (*ADRA1A, ADRA1B, ADRA1D, ADRB1, ADRB2, ADRB3*) and from the cholinergic group (*CHRNA4, CHRNA7, CHRM3*) were also quantified. *ADRA1A, ADRB2* and *CHRNA4* were the most highly expressed. Transcript abundance is shown for both groups in **Suppl. Figure 10**. Data indicate the regulation of several receptor transcripts in tumors, using peritumoral (F4) areas as our reference. Within the adrenergic group, *ADRA1A* and *ADRB1* displayed a down-regulation of more than 2-fold, whereas *ADRA1D* and *ADRB2* were up-regulated by 3.2-fold and 2-fold, respectively. Within the cholinergic group of transcripts, the nicotinic receptor mRNA *CHRNA4* was profoundly depleted (> 10-fold) in HCC versus F4 samples, while the muscarinic receptor *CHRM3* was up-regulated by 3-fold. Modulated transcripts *ADRA1A, ADRB2* and *CHRNA4* were also the most highly expressed (> 1,000-fold in comparison with the others) and therefore also likely participated in ANS-regulated phenotypes concomitant to the F4-to-HCC transition. Altogether, such data suggest ANS-driven changes to parenchymal signaling during the F4 to HCC transition. Regulation of these receptors confirms at the post-synaptic level the remodeling of upstream ANS neurons, as identified in other types of cancers (4–8).

### Tumor-specific uncoupling between neuronal and post-synaptic orientations

In order to more closely monitor the potential sensitivity of neural processes to tumorigenesis, we next sought to identify concordant or conflicting neural orientations at the pre- and post-synaptic levels in non-tumoral versus tumoral tissues. Again, neural TH and VAChT were quantified by WB. At the post-synaptic level, the three most highly expressed transcripts were used (*ADRA1A*, *ADRB1, ADRB2* for the adrenergic polarity and *CHRNA4, CHRNA7, CHRM3* for the cholinergic polarity). In F4 peritumoral samples, shifts of ratios defined by adrenergic over cholinergic signals remained homogenous across paired specimens when comparing neural markers and post-synaptic markers (p < 0.0001). In stark contrast, in HCC, shifts were unpredictable across samples, and even contradictory with respect to F4 features (**Figure 2B-C** and **Suppl. Figure 11**). Altogether, such data indicate that hepatic tumorigenesis is correlated with the empowerment of the tumor, ignoring neural polarities observed in the non-tumoral, cirrhotic, tissue.

### Bioinformatics illuminates the pathogenic implication of the parasympathetic orientation in HCC evolution

#### General sample characterization

We then sought to decipher the potential implication of the ANS in HCC phenotypes and stratification, through bioinformatics. In order to charter the interplay between autonomic functions and HCC in the least possibly biased manner, we performed a multisectional bioinformatics study on the previously published HCC TCGA dataset (19).

The adrenergic/cholinergic signal balance defines a unified ANS output in each innervated organ. We first defined a signature encompassing adrenergic and cholinergic receptor transcripts considering all mRNAs belonging to both categories. All these receptors (n = 24) are listed in the **Suppl. Table 3.** Adrenergic and cholinergic gene set normalized scores were established and the difference between adrenergic and cholinergic scores was then calculated. Based on the resulting score, samples were split into two classes: those with a higher difference than median were named adrenergic and those with a lower difference than median were named cholinergic (see the Materials and methods section). The mathematical formula used, distribution diagram of obtained values and PCA representation of those two classes, thereafter named adrenergic or cholinergic classes, define two populations and are shown in **Figure 3A-C.** We then generated data describing normalized expression levels of all adrenergic and cholinergic receptors in HCC, in all TCGA samples, at the level of specific etiologies and tumor differentiation statuses. Differential expression analysis revealed that *ADRA1A, ADRA1B, ADRA1D, ADRB1, ADRB2, CHRM1, CHRM4, CHRNA1, CHRNA3, CHRNA5, CHRNA6, CHRNA7, CHRNA9, CHRNB2, CHRNB4* were expressed at various levels in both neural classes. In general, as expected, ADRAx and ADRBx receptors were better expressed by the adrenergic class, while CHRNAx and CHRNBx receptors were better expressed in the cholinergic class. (**Figure 3D** and **Suppl. Table 3**).

**Figure 3.**
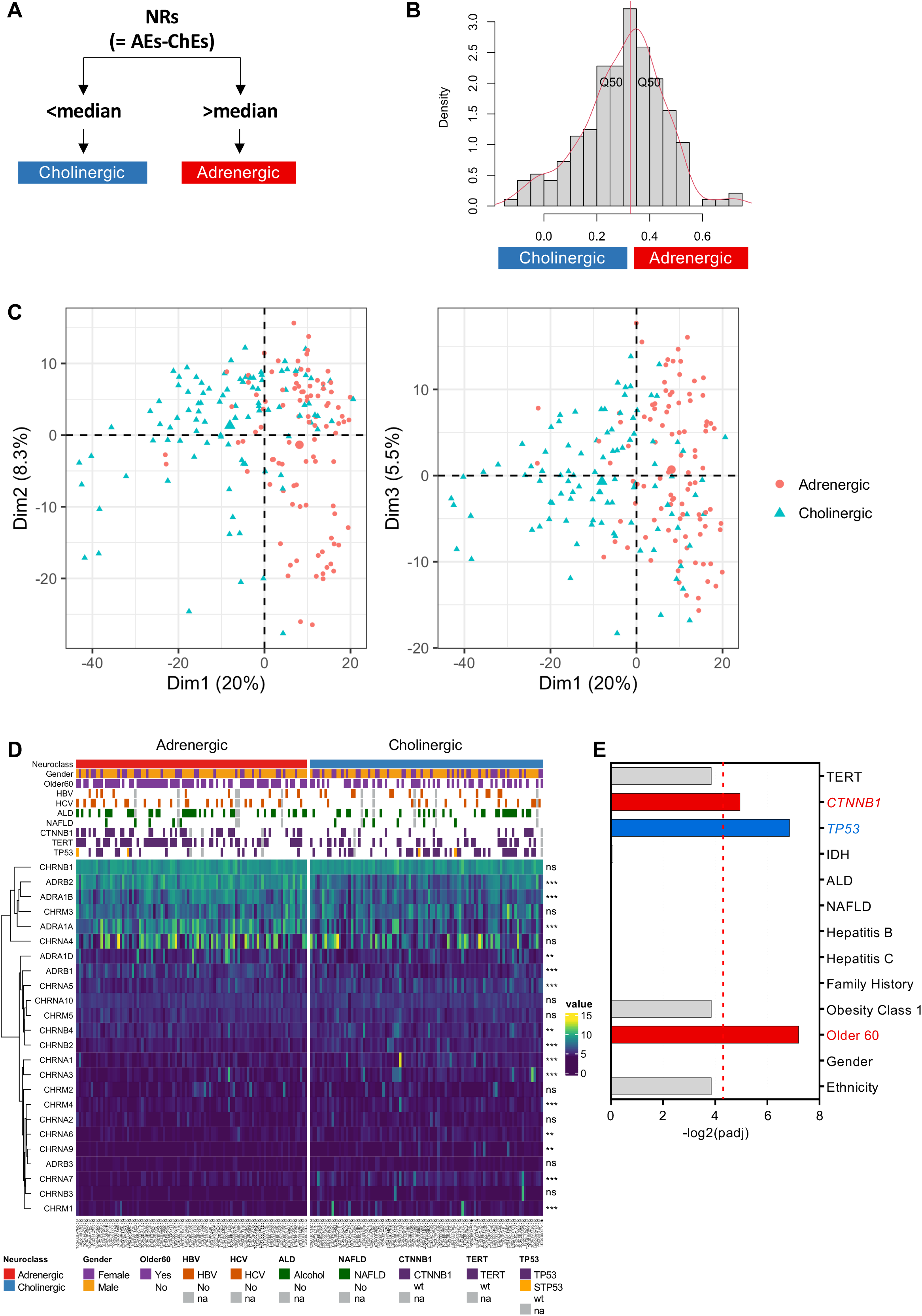
Neurosignature based on adrenergic and cholinergic receptors in HCC. **A.** Neuronal receptor score (NRs) calculation: adrenergic and cholinergic enrichment scores (AEs and ChEs, respectively) were calculated based on corresponding receptors gene set signatures for each sample, then cholinergic scores were subtracted from adrenergic scores. This simple approach enabled process of normally distributed values. Samples were grouped in cholinergic and adrenergic classes depending on whether their ssGSEA score values (NRs) were below or above the calculated median, respectively. **B.** Histogram of NRs calculated by ssGSEA for HCC tumors. **C.** Sample distribution after dimensional reduction (PCA) based on the 10 per cent most variable genes (normalized values). Cholinergic class in blue and adrenergic class in red are contrasted. **D.** Heatmap of relative expression levels of all adrenergic and cholinergic receptors in HCC tumor samples. Once samples were classified in cholinergic and adrenergic receptor expression classes based on previously obtained signature scores, differential expression analysis was performed and expression values normalized (variance stabilization transformation) and shown in the heatmap. Older60: patients older than 60. HBV: Hepatitis B Virus; HCV: Hepatitis C Virus; ALD: Alcoholic Liver Disease; NAFLD: non-alcoholic fatty liver disease. Mutations: TP53, CTNNB1, TERT; wt: wild type. na: no information available. **E**. Associations between classes and main HCC clinico-biological features. Fisher test’s adjusted *p*-values per variable (** p < 0.01, *** p < 0.001).

#### Association between neural orientations and TP53 and CTNNB1 mutations

In order to identify a potential association between a given ANS polarity (sympathetic or parasympathetic) and classically considered clinico-biological parameters in the HCC field, we implemented two-dimensional PCA on these data with respect to sex, ethnicity, etiology, obesity and mutational profile (*hTERT, TP53, CTNNB1*) (**Suppl. Fig. 12**). Importantly, no association could be seen between either neural class and sex, ethnicity, obesity, family history, or any of the four main HCC etiologies (HBV, HCV, NASH, ALD). *CTNNB1* mutations and age >60 emerged as positively associated with the adrenergic class. *TP53* mutations emerged as positively associated with the cholinergic class (padj ≤ 0.05). Of note, mutated *CTNNB1* samples positively correlated with the adrenergic class, yet at the significance threshold of 0.05 (**Figure 3E**). A Fisher’s exact test comparing each class signature to each variable was constructed (**Suppl. Table 4**). This suggests that the adrenergic and cholinergic classes defined by this ANS-based signature may enrich the stratification of HCC heterogeneity, this time based on mutual influences or selective events occurring between intrahepatic neural inputs and canonical HCC genetics.

#### Differential Gene Expression and Pathway Enrichment Analysis

To identify which HCC phenotypes clustered with the ANS features of the tumor, we performed a differential gene expression analysis (padj < 0.01 and absolute Log2 fold-change > 0.58), the results of which are illustrated in a Volcano plot (**Figure 4**). Data related to the 100 most significantly up- and down-regulated genes for the adrenergic and cholinergic signatures are shown in the **Suppl. Tables 5-6,** respectively. Several mRNAs up-regulated in the cholinergic signature are of pathological significance: (i) *LGALS14* transcript (8-fold Log_2_), which encodes a protein fostering apoptosis of T cells, (ii) *CT55* (cancer testis antigen, ranked 3^rd^; +6-fold Log_2_) (iii) *BMP7*, ranked 5^th^ (+6-fold Log_2_), is a growth factor of the TGF-β family, *CEACAM7* (carcinoembryonic antigen-related cell adhesion molecule 7, ranked 7^th^; +5-fold Log_2_) and *MAGEA4* as well as *MAGEA10* (ranked 10^th^ and 30^th^; +4-fold Log_2_ each; both related to MAGEA3, a novel HCC progression driver), XAGE2 (ranked 44^th^; +4-fold Log_2_), a fetal / reproductive tissue tumor-related antigen, and GAGE2A, (76^th^; +3-fold Log_2_) a germ-cell and tumor antigen. CEA, MAGE, XAGE and GAGE antigens are related to colorectal cancer, melanoma and fetal/reproductive tissue tumors, respectively, suggesting a link between cholinergic orientation and general cancer markers. Likewise, *AFP* (+4-fold), ranked 33^rd^ in terms of positive association with the cholinergic signature, encodes a widely used protein for HCC diagnosis, as well as tumor size and differentiation evaluation (**Suppl. Table 5**). In contrast, one alcohol dehydrogenase (ALDH3A1; +3-fold) and three major CYP450 isoforms (1A1, 3A4, 1A2), as well as CYP2A13, 2A7P1 and 3F36P, several of which known to be xenobiotic-inducible hepatocytic differentiation markers, were enriched (2 to 3-fold Log_2_) within the transcripts significantly associated with the adrenergic class (**Suppl. Table 6).** The β-catenin control target *GLUL*, but not *LGR5*, was also positively associated with this adrenergic class (+1.52 Log_2_; padj = 4.6E-8, rank 241). In contrast, no CYP450 mRNA was found in the cholinergic class. Altogether, such data support that the cholinergic signature may be correlated with less differentiated HCC tumors.

**Figure 4.**
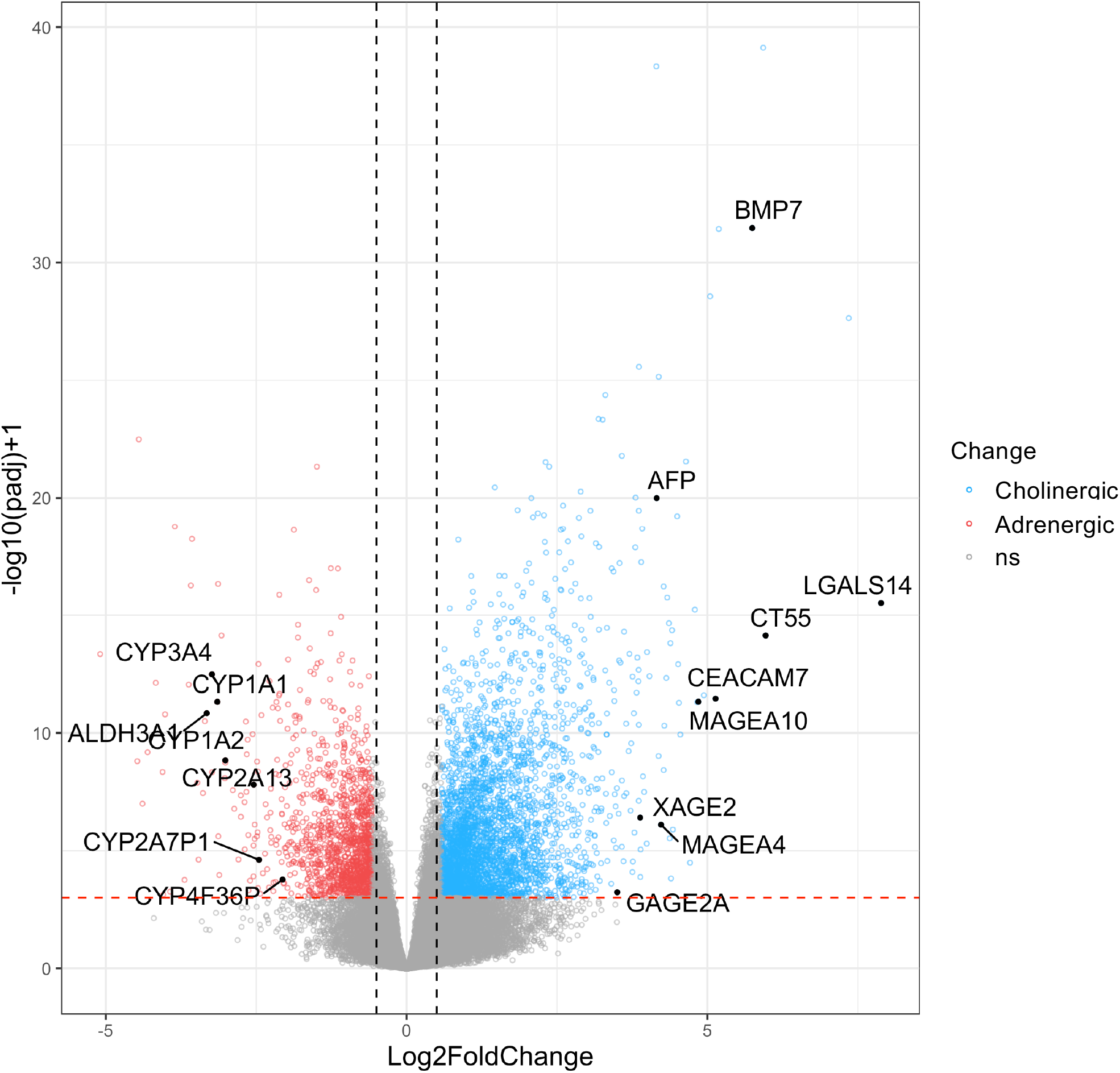
Transcriptomic features of adrenergic and cholinergic signatures analyzed by DGE. Volcano plot showing differentially-expressed genes in both tumor classes. Genes with lowest adjusted p-value are indicated. The horizontal red bar represents the adjusted p-value threshold at 0.01. The higher the location of a gene, the stronger its association with a low adjusted p-value (significance). The more a gene is located to the right or left, the stronger its association with a high absolute Log2 fold-change. Modulated genes were considered if absolute Log2 fold-change value was higher than 0.58 and adjusted p-value < than 0.01.

#### Pathways associated with ANS polarities by overrepresentation analysis

Next, to accurately document how ANS features of the tumor are integrated within the current landscape of signatures in HCC, we performed an overrepresentation analysis (21) for both adrenergic and cholinergic classes using ClusterProfiler (24) and the msigdbr R package (23) to access Molecular Signature Databases for gene sets H, C2, and C5. The lists of genes obtained from differential gene expression were used as inputs. A total of 956 and 604 pathways were correlated with adrenergic and cholinergic signaling, respectively (padj < 0.01, **Suppl. Tables 7-8**). We first sought to validate the approach by probing the presence of known recognized adrenergic and cholinergic processes in both cases. Overrepresentation analysis of the adrenergic class identified 24 pathways related to cardiac or heart processes, mainly sympathetically regulated, and none related to nausea or vomiting, hallmarks of parasympathetic activation. Within the analysis pertaining to the cholinergic orientation, two signatures were related to nausea and vomiting, triggered by parasympathetic activation, while none were associated with cardiac pathophysiology.

We then deciphered the most relevant differences between overrepresentation pathways of adrenergic and cholinergic tumors. Pathways shared by all cancers (chosen keywords were ‘cancer’ and ‘tumor’) were more associated with the adrenergic class (82 (57%) pathways for adrenergic tumors vs 63 (43%) for cholinergic tumors), (**Figure 5A-B**). In contrast, HCC-specific pathways were enriched in the cholinergic class of tumors (70% of the total number of HCC pathways identified herein versus 30% in the adrenergic group). Significance of the pathways was also higher in the cholinergic class, as illustrated by the frequency diagram (**Figure 5C-D**). In the case of adrenergic signaling in HCC, a stronger association was also found with immunological functions (72% vs 28%, **Figure 5E-F**). Strikingly again, 97% of energetic pathways, representing functions fostering general proliferation, were correlated to cholinergic tumors (**Figure 5G-H**), whereas, as seen in overrepresentation pathways data, specific pathways linked to *CTNNB1* mutations were associated with the adrenergic tumor class.

**Figure 5.**
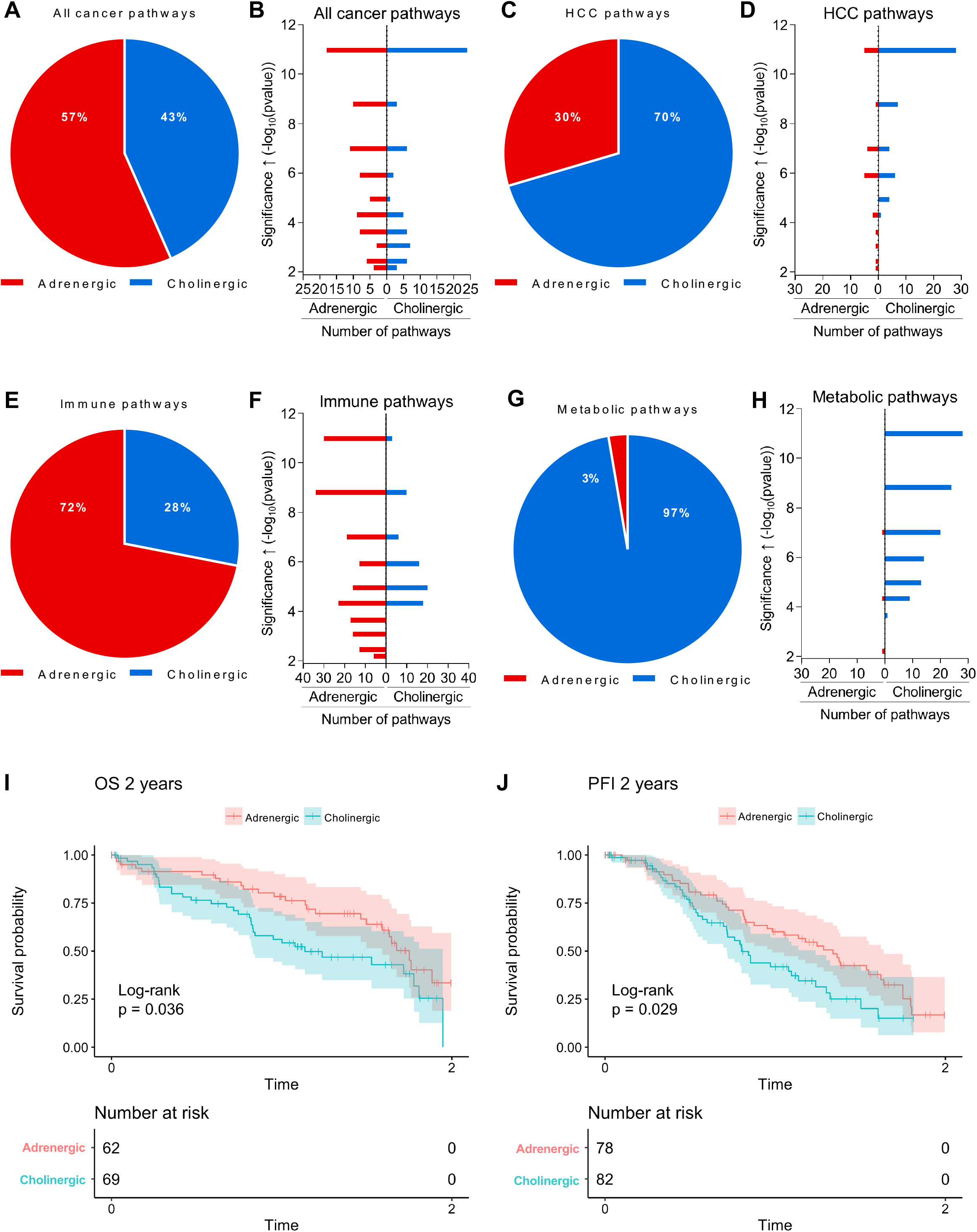
Cholinergic orientation correlates with adverse HCC pathways and outcomes as identified by Gene Set Enrichment Analysis (GSEA). From up-regulated (derived from adrenergic tumors) and down-regulated (derived from cholinergic tumors) transcripts obtained by differential gene expression analysis, enriched pathways were identified. Pathways with adjusted p-value < 0.01 were selected. **A,C,E,G**. Pathway allocation to each neuroclass. **B,D,F,H**. Statistical significance of both groups of pathways. The higher the pathway on the plot (low *p*-value, then (-Log)-transformed), the stronger its association with the neuroclass of interest. Key-words used for the search are: Cancer terms: CANCER|TUMOR. HCC terms: HEPATOCELLULAR_CARCINOMA|HCC Immune terms: IMMUN|MACROPHAGE|PHAGO|IGM|ANTIGEN|LEUKO|CYTO|TREG|LYMPHOCYTE|CHE MOKINE|INTERLEUKIN|MONOCYTE|T_CELL|NEUTROPHIL|MACROPHAGE|CD8|CD4|REGULATORY_T|DENDRITIC|B_CELL|NK_CELL|NATURAL_KILLER|INFECTION|INFLAMATI ON. **I-J.** Kaplan-Meier representation of the predictive value of both classes with respect to overall survival (OS) and progression-free interval (PFI) in HCC.

We finally sought to identify within both neural classes the recognized HCC-specific signatures known to be related to good or poor prognosis pathways, including those identified by Chiang (34), Hoshida (35), Lee (36) and Boyault (37) that are regularly considered for classification (2). The adrenergic class was associated with 18 known signatures, 17 of which were in functional agreement with the transcriptomics of this class, i.e. related to better prognosis. The cholinergic class was linked to 8 signatures, of which 7 were in agreement with the transcriptomics of this class, that is, related to poor prognosis (**Suppl. Table 9**). In addition, the adrenergic class was correlated with longer overall survival (OS) and progression-free interval (PFI) than its cholinergic counterpart within a timeframe of two years post-diagnosis when performing a Cox model (**Figure 5I-J** and **Suppl. Table 10**), corroborating our findings that the cholinergic class is associated with more aggressive tumors.

To interpret these data at the level of multigenic functions, we performed a single sample non-parametric, unsupervised gene set variation analysis (GSVA method ssGSEA) for each sample, using the *Hallmarks of Cancer* signatures (38). As shown in **Figure 6A**, the adrenergic class was correlated with *CTNNB1* mutation-associated metabolic functions, such as oxidative phosphorylation, adipogenesis, peroxisomal activity, xenobiotic detoxication, bile acid functions, and fatty acid metabolism (padj < 0.05). In contrast, the cholinergic class showed a clear association with mitogenic processes, E2F, MYC, TGF-β signaling, EMT, angiogenesis, NOTCH, Hedgehog, KRAS, TNF-α/NF-KB, IL6/STAT3, P53, hypoxia and PI3K/Akt/mTOR signaling pathways (padj < 0.05) (**Suppl. Table 11**). Hence, adrenergic signaling was correlated with several metabolic functions defining the non-proliferative class, whereas cholinergic signaling was correlated with features defining the proliferative class of HCC tumors (3). Again, these findings argue in favor of a poorer evolution for HCC patients with higher cholinergic signaling.

**Figure 6.**
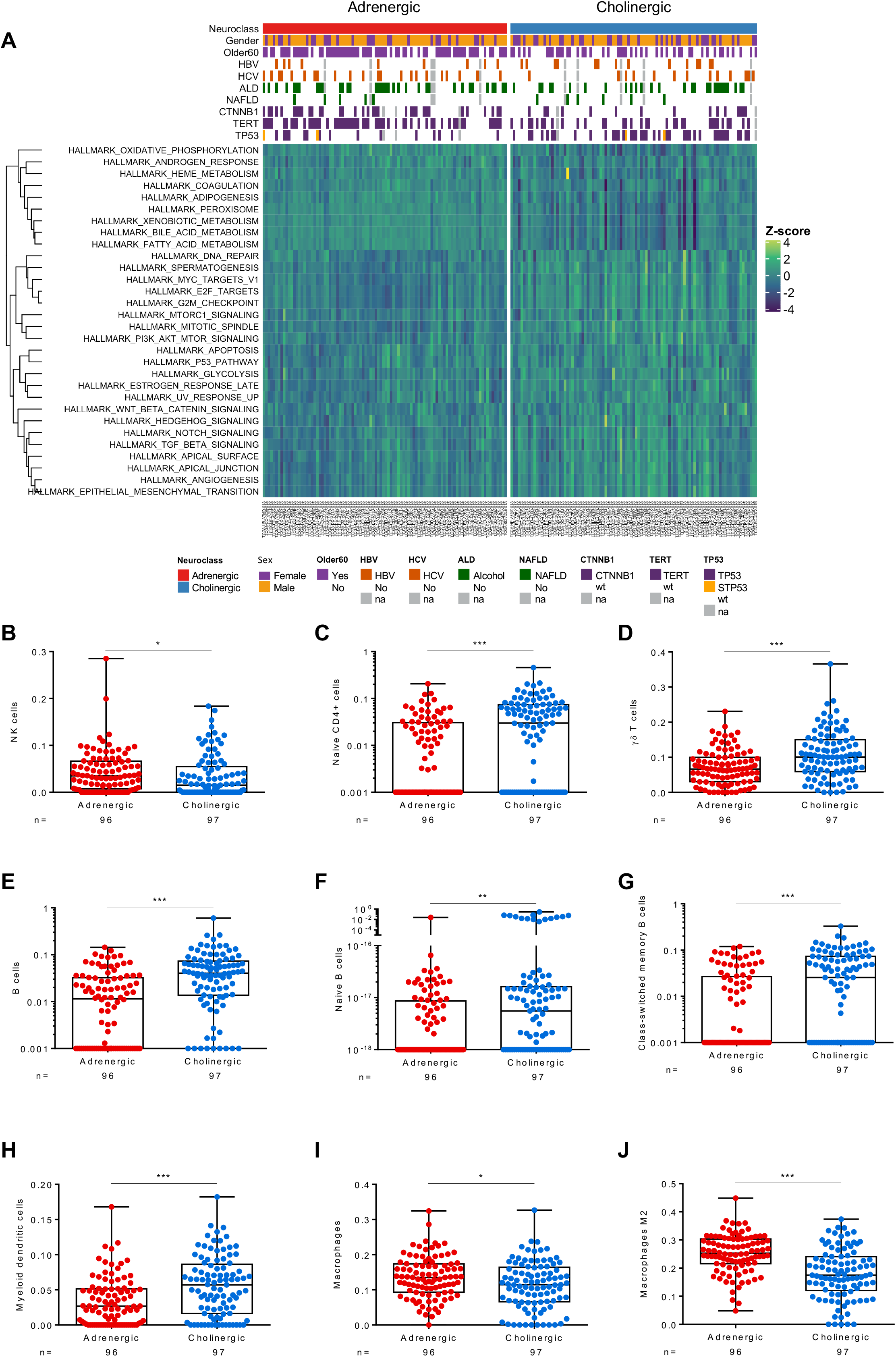
Adrenergic- and cholinergic-associated hallmark pathways determined by single sample Gene Set Enrichment Analysis (ssGSEA), and comparative immune infiltrations determined by the xCell method. **A**. ssGSEA scores were calculated for each sample. Pathways comparison between neural classes was performed with the Wilcoxon Test and significant pathways (padj < 0.05) selected. Older60: patients older than 60. HBV: Hepatitis B Virus; HCV: Hepatitis C Virus; ALD: Alcoholic Liver Disease; NAFLD: non-alcoholic fatty liver disease. Mutations: TP53, CTNNB1, TERT; wt wild type. na: no information available. To determine statistical differences for each cell type between HCC neural classes, a Wilcoxon Test was performed on 36 immune subpopulations using the immune Xcell score. Significant data obtained on 9 of them are shown (panels **B-J**, padj < 0.05).

#### Correlation between ANS orientation and immune response

Having observed that the number of immune pathways linked to adrenergic tumors was higher than to cholinergic tumors, and given the known impact of HCC immunology on clinical outcome, we then considered potential associations between ANS features and intra-tumoral immune markers, deciphered using the xCell enrichment method (30). Interestingly, neither global immune infiltrates nor T CD8+ leucocytic populations were correlated with either class. However, NK cells (+1.3-fold) were enriched in adrenergic tumors. Instead, CD4+ T cells (naïve, +2.5-fold), gamma/delta T cells (+1.5-fold) and several types of B cells (composed of total (+2.7-fold), naive (+29-fold) and class-switched (+2.3-fold) subpopulations) as well as myeloid dendritic cells (+1.7-fold) were significantly enriched in the cholinergic class. Total and M2 macrophages were somewhat unexpectedly enriched (+1.2 and +1.4-fold respectively) in the adrenergic class (padj < 0.05) (**Figure 6B and Suppl. Table 12).** Overall, such data suggest that a B cell orientation, known to foster immunotolerance, is associated with the cholinergic polarity of the tumor. Vagal cholinergic signaling conveys systemic anti-inflammatory signaling (39) as well as in the gastrointestinal tract (40, 41). These findings identify cholinergic signaling as a potential contributor or intrinsic signature of specific pro-tumoral or exhausted immune subsets, likely implicating B cells, which are newly identified and debated players in HCC (42–45). Accordingly, we investigated the association between each class and current immunotherapeutic targets of HCC. A hit was found linking one altered PD1 levels-associated pathway and the adrenergic class (GSE26495_PD1HIGH_VS_PD1LOW_CD8_TCELL_UP, padj = 3.5E-3, see **Suppl. Table 7**). This indicates that the transcriptome of adrenergic tumors is enriched in specific genes that are themselves correlating with variations of PD1 levels. Overall, these data suggest that ANS polarities are correlated with immune processes that are known to influence pathology progression.

## Discussion

The involvement of the ANS in the pathophysiology of many diseases is now recognized and links the CNS, which is unique to each patient, to disease-specific peripheral processes. HCC, a highly heterogeneous cancer at the genetic and pathological levels, remained undocumented with respect to the participation of the ANS. Local neo-neurogenesis and the involvement of the ANS has been identified in ovarian (4), prostate (5), gastric (6) and pancreatic (7, 8) cancers. In prostate cancer, a study revealed cancer-associated neo-neurogenesis with DCX marker expression, and predominant parasympathetic (cholinergic) features of *de novo* grown intra-tumoral nerves (46). In gastric and pancreatic cancers, vagal nerves regulate cholinergic receptors in favor of oncogenesis initiation and progression. In breast cancers, in contrast, a study showed that parasympathetic nerves reduced cancer growth and progression (47). In the case of HCC, DCX/VAChT staining immune localization, as well as Western blots advocate for an association between the parasympathetic features of the liver ANS and the disease. As was shown recently in NASH (33), sympathetic innervation of HCC seems to be weaker if existent. Drastic enrichment of neurogenetic pathways in the adrenergic class identified in our work could thus be interpreted as compensatory with the aim of reestablishing sympathetic neural control in the neuro-emancipated diseased tissue. Indeed, it would be interesting to know whether HCC-correlating neurogenic netrin-1 (48) as well as NGF and neurotrophin 5 (increased in NASH (49)) are associated with the adrenergic class.

Herein, we identify parasympathetic innervation as dominant over sympathetic innervation in human, as well as in cirrhotic rat HCC. Weakening of cognitive functions can be observed in elderly patients under medication with anti-cholinergic drugs. In line with two previous studies {Shaked, 2009 #7356;Hanin, 2018 #7359}, the increase in the cholinergic intrahepatic system that correlates with disease progression may also derivate from a central attempt to dampen deleterious, liver-related, brain effects. In a more HCC specific manner In addition, we demonstrate that cholinergic tumors are associated with higher *AFP* transcript levels and many poor prognosis-related pathways. Immune tumor infiltrates are correlated with better prognosis in HCC (50, 51). The cholinergic class also seems to be partly depleted of cytotoxic immune or immunotherapy-related markers, suggesting their partially weakened or exhausted immune status. The liver is innervated by vagal outputs, which bear anti-inflammatory properties in several instances including the gastrointestinal tract (52). This could provide an explanation for higher aggressiveness of cholinergic tumors, as cholinergic/vagal mitigation of local immunity, as suggested by the data herein, could obstruct proper mounting of anti-tumoral responses. In average, *CTNNB1*-mutated specimens that correlate with adrenergic polarity herein are in fact known to be rather immune-excluded (2). This is suggesting that a somewhat heterogenous group of adrenergic tumor subtypes, not yet stratified, may account for the observed association between *CTNNB1*^mut^ and adrenergic tumors. Indeed, this association only reaches the null hypothesis threshold of 0.05 and implicates the *GLUL* control transcripts enrichment but not the *LGR5* control transcript enrichment, suggesting its moderate significance. Most importantly, our data both further validate and enrich the current landscape of predictive transcriptomic signatures, showing, once again the intensified association between adrenergic features and a series of transcriptomic signatures linked to better prognosis.

The novelty of our findings resides in the implication that the systemic regulation of each patient influences the features of their tumor. In the liver, the *fight-or-flight* model, that is generally accepted for describing ANS functions (53), would predict that adrenergic signaling would mobilize intracellular hepatocytic energy pools for peripheral energetic needs, whereas cholinergic signaling would foster intrahepatic storage of nutrients and related processes, such as liver expansion. This model seems pertinent to the HCC context. Liver expansion implicates liver cell size/mass increase as a condition for organ growth, both being dependent on mTOR functions (54, 55), a pathway that is correlated with cholinergic tumors in our study. Frequent comorbidities associated with liver carcinogenesis are excessive BMI and alcohol intake (an underestimated but important energetic source), suggesting that parasympathetic signals aiming at fostering liver expansion, due to their implication in the rest-and-digest related functions {LeBouef, 2022 #7357}, could be highjacked by the tumor in a context of excessive nutrient availability. This is in accordance with the so-called antagonistic pleiotropy hypothesis {Rose, 1980 #7360}, that indicates that genes that enhance fitness early in life may diminish it in later life. The likely protective and adverse roles of coffee (56) and tobacco (56, 57), respectively, as adrenergic and cholinergic agonists, are also in support of our findings.

This study provides the first body of evidence with respect to neural implication in human HCCs and in their stratification, using ANS functions unique to each patient’s pathophysiology.

## Acknowledgements

Non-TCGA clinical samples were used thanks to the participation of the French Liver Biobanks network (INCa, BB-0033-00085).

## Supplementary Figures

**Supplementary Figure 1.**

Histological assessment of disease progression in the DEN HCC rat model. Hematoxylin/Eosin and Sirius Red staining were used.

**Supplementary Figure 2.**

**Validation of anti-NeuN, DCX, INA, TH and VAChT antibodies by Western Blot.**

**A.** Extracts from neonate mouse brain, SKNSH, PHH, and human HCC lines belonging to transcriptomic classes 1, 2 and 3 (58) were processed for detection of the indicated targets.

**B.** The same strategy was used for the validation of an anti-DCX antibody specifically suitable for rat epitopes. **C-E.** Signal quantification of data presented in **Suppl. Fig. 3-7** was done using the Fiji software on non-saturated images, prior to statistical plotting using the Mann-Whitney test (* p < 0.05). Quantifications conducted prior to (B-C) and after (D-G) patient segregation according to the differentiation grade of the disease. HCC: Hepatocellular carcinoma; HCV: Hepatitis C Virus; HBV: Hepatitis B Virus; ALD: Alcoholic Liver Disease; NAFLD: non-alcoholic fatty liver disease. Peritumoral: cirrhotic; Tumoral: HCC; HCC differentiation grade: Low/moderate or High.

**Supplementary Figure 3.**

**Quantification of mature and progenitor neural markers in human HCC of all etiologies.**

**A-L.** Signal quantification was done using the Fiji software on non-saturated images prior to statistical plotting using the Mann-Whitney test (n = 51 patients, n.s). Quantification before (A-F) and after (G-L) patient stratification based on the ‘low/moderate’ or ‘well’ level of differentiation.

**Supplementary Figure 4.**

**Expression of mature and progenitor neural markers in human HCC of HBV etiology.**

**A.** Immunoblotting on neural markers using antibodies against NeuN, DCX, TH and VAChT neural markers and *β*-tubulin as an internal control. **B-G.** Signal quantification was done using the Fiji software on non-saturated images prior to statistical plotting using the Mann-Whitney test (n = 14 patients, * p < 0.05, ** p < 0.01, *** p < 0.001).

**Supplementary Figure 5**.

**Expression of mature and progenitor neural markers in human HCC of HCV etiology.**

**A.** Immunoblotting on neural markers using the above-mentioned antibodies. **B-G.** Signal quantification was done using the Fiji software on non-saturated images, prior to statistical plotting using the Mann-Whitney test (n = 9 patients, * p < 0.05, ** p < 0.01, *** p < 0.001).

**Supplementary Figure 6.**

**Expression of mature and progenitor neural markers in human HCC of NASH etiology. A.** Immunoblotting on neural markers using the above-mentioned antibodies. **B-G.** Signal quantification was done using the Fiji software on non-saturated images, prior to statistical plotting using the Mann-Whitney test (n = 14 patients, * p < 0.05, ** p < 0.01, *** p < 0.001).

**Supplementary Figure 7.**

**Expression of mature and progenitor neural markers in human HCC of ALD etiology.**

**A.** Immunoblotting on neural markers using the above-mentioned antibodies. **B-G.** Signal quantification was done using the Fiji software on non-saturated images, prior to statistical plotting using the Mann-Whitney test (n = 14 patients, * p < 0.05, ** p < 0.01, *** p < 0.001).

**Supplementary Figure 8.**

**Validation of antibodies used by IHC**. Anti-NeuN, DCX, TH and VAChT antibodies were validated by immunofluorescence on ad hoc human tissue samples prior to their use on HCC specimens. **A.** Cerebellum. **C.** Parotid gland tumor. **E.** Surrenal gland. **G.** Cerebral cortex.

**B, D,F,H.** Normal liver (hepatocytic areas). Nuclei were stained with DAPI (grey signal). Antigens of interest are shown in green. Scale bar = 100 μm.

**Supplementary Figure 9.**

**Comparison of post-synaptic receptors mRNA expression levels upon comparison with non-tumoral paired tissues.**

Adrenergic and cholinergic receptors abundance and expression in peritumoral tissue (F4) and HCC. 166 pairs of liver tissues were analyzed. Mann-Whitney test (* p < 0.05, ** p < 0.01, *** p < 0.001).

**Supplementary Figure 10.**

**Human HCCs harbor DCX+ neurogenesis with predominant parasympathetic features.** A panel of 24 HCC samples derived from all four main etiologies (HBV: n = 7; HCV: n = 4; ALD: n = 9; NASH: n = 4) was probed for immunolocalization of NeuN, DCX, TH and VAChT by multiplex IHC. **A-F**. Masson’s trichrome and IHC staining of a representative capsule-bearing tumor. **G-J.** Staining of a representative tumoral bulk. Nuclei appear as grey. Cap: capsule; Tu: tumoral bulk. Scale bars = 50 μm.

**Supplementary Figure 11.**

**Relationship between neuron marker and post-synaptic receptors expression after etiology-based stratification.**

TH and VAChT expression levels in peri-tumoral and tumoral samples (n = 51 HCC cases) were evaluated by immunoblotting with specific antibodies and normalized against whole lane signal. Gene expression levels of adrenergic and cholinergic receptors were evaluated by qPCR and normalized against *GUS* expression. Adrenergic/cholinergic ratio was calculated for each patient as follows: (ADRA1A+ADRB2+ADRA1B) / (CHRNA4+CHRNA7+CHRM3), where those six targets are the most expressed of both categories in patient samples. Mann-Whitney test (* p < 0.05, ** p < 0.01, *** p < 0.001).

**Supplementary Figure 12.**

**HCC neural classes are unrelated to many HCC clinico-biological parameters except age>60, *CTNNB1* and *TP53* mutational status.** PCAs comparing signature classes with sex, ethnicity, obesity, family history, or any of the four main HCC etiologies (HBV, HCV, NASH, ALD), or main mutational profiles *IDH, TP53, TERT, CTNNB1* mutations.

## Supplementary Tables

**Supplementary Table 1.**

Characteristics of paired F4/HCC samples used in the study.

**Supplementary Table 2.**

Significance assessment of neural features in HCC.

**Supplementary Table 3.**

Differential gene expression (by DESeq2) of adrenergic and cholinergic receptors in TCGA samples grouped by neural class.

**Supplementary Table 4.**

Association levels between neural classes and major HCC-related clinico-biological variables. Fisher test. p<0.01.

**Supplementary Table 5.**

List of the 100 genes most correlated with the cholinergic class.

**Supplementary Table 6.**

List of the 100 genes most correlated with the adrenergic class.

**Supplementary Table 7.**

Pathways enriched in the adrenergic class. Over Representation Analysis with hypergeometric test. p < 0.01.

**Supplementary Table 8.**

Pathways enriched in the cholinergic class. Over Representation Analysis with hypergeometric test. p < 0.01.

**Supplementary Table 9**.

ssGSEA HCC signatures correlated with adrenergic (left columns) and cholinergic (right columns) classes of tumors. Pathways are listed by decreasing order of significance. Besides signature names listed in this table (p < 0.01 Wilcoxon Test), attention needs to be paid to their expanded biological significance extracted from the ssGSEA database, listed **in Suppl. Table 9**, before drawing conclusions with respect to pathology.

**Supplementary Table 10.**

Biological descriptions of prognosis pathways associated with both HCC neural classes.

**Supplementary Table 11.**

Significance of hallmark pathways. Wilcoxon Test p < 0.05.

**Supplementary Table 12.**

Significance of immune alterations. Wilcoxon Test p < 0.05.

**Supplementary Table 13.**

Antibodies used in this study.

**Supplementary Table 14.**

Human qPCR primers.

