## Supplementary material for "Hepatocellular carcinoma hosts immature neurons and cholinergic tumors that correlate with adverse molecular features and outcomes": Suppl Info 1

**Supplementary Information 1**

**Supplementary methods**

**Rat liver samples**

Methods conform to the ARRIVE guidelines. 6-week-old Fischer 334 male rats (Janvier Laboratories) were acclimatized for two weeks, given DEN at 50 mg/kg weekly from day 0 to week 14, allowing progression of chronic liver disease to fibrosis, cirrhosis, decompensated cirrhosis and HCC. The experimental unit was a cage of 3 to 4 animals for a total of 28 animals that were fed *ad libitum*. One to two experimental units were allocated to each group for a total of 28 rats. Five rats represented the untreated group. Seven rats represented the DEN+ fibrotic group. Eight rats represented the DEN+ cirrhotic group, likewise in the DEN+ HCC group. Previous publications on this model indicate that at least 7 rats are necessary in DEN-treated groups for robust statistics. No rat was excluded from the analysis. Mixing of all groups in similar proportions was used to allocate experimental units to control and each treatment groups. Animals were sedated using ketamine prior to sacrifice. The outcomes measures assessed are the molecular markers depicted in **Figure 1**. Spearman and Mann-Whitney tests were used under GraphPad/Prism. The status of the data related to the null hypothesis was considered. Rat samples were obtained under the Grenoble-Alpes University agreement #B 38 516 10 006. No conflict of interest interfered with the study’s design.

**Western blotting**

Immunoblotting was performed using 40 μg of RIPA (50 mM Tris HCl (pH 8.0), 150 mM NaCl, 1% NP-40, 0.5% sodium deoxycholate, 0.1% SDS, 10 mM sodium fluoride, 50 mM orthovanadate, 1x protease inhibitor cocktail (Roche))-processed cell lysates, then resolved on 8% or 10% SDS-PAGE, blotted onto nitrocellulose membranes (Amersham Biosciences, Saclay, France), blocked using 5% low fat dried milk in TBS Tween 0.1% for 1 h at room temperature (RT) and probed overnight at 4°C with corresponding antibodies listed in the **Suppl. Table 13**. After three washes in TBS-Tween 0.1%, membranes were incubated for 1 h at RT with secondary antibodies coupled to HRP (1/5,000, Sigma-Aldrich, St-Quentin, France) prior to chemiluminescence-based visualization using the Clarity Western ECL substrate (Bio-Rad, Versailles, France). Ponceau S fluorescence levels were used for normalization because of high variability of housekeeping protein signals in clinical samples, as verified in the present study (**Suppl. Figures 4 to 7**).

**Total RNA extraction and RT-qPCR**

Total RNA was extracted using Trizol (Invitrogen). One μg of RNA was DNAse I-digested (Promega, Charbonnières, France) and reverse transcribed using SuperScript VILO reverse transcriptase (ThermoFischer, Les Ulis, France) according to the manufacturer’s instructions. Quantitative real-time PCR was performed on 1/5^th^ diluted samples on a LightCycler 96 device (Roche, Meylan, France) using the No Rox qPCR mix (Bioline, Paris, France). PCR primer sequences (5′-3′) and qPCR conditions are listed in **Suppl. Table 13**. Specificity of all primers was assessed by melting curve analyses and agarose gel electrophoresis. Efficiency of all primers was quantified using 3-fold serial dilutions of target templates.

**Immunohistochemistry**

All reagents were from Sigma-Aldrich (St. Quentin Fallavier, France) unless otherwise stated. Samples where fixed for 24 h in 4% formaldehyde (pH 7.0). Progressive dehydration was performed using ethanol and xylene through 70%, 80%, and 95% ethanol (45 min each), followed by 3 changes of 100% ethanol (1 h each). Tissue was cleared through 2 changes of xylene, for 1 h each. Tissue was then immersed in 3 changes of paraffin, for 1 h each. Inclusion was automatically performed in a Histocentre3 Shandon (Loughborough, United Kingdom) device prior to cutting 4 µm-thick sections, transferred to SuperFrost slides (Euromedex, Souffelweyersheim, France), and dehydrated at 56°C for 1 h. For Masson’s trichrome staining, the whole procedure was performed in a Leica ST5020 (Jena, Germany) apparatus. Paraffin removal was done using xylene and ethanol (backwards compared to protocol depicted above), followed by water rinsing. Gill hematoxylin was then added for 10 min followed by water rinsing. Saturated lithium carbonate was then used for 5 s before rinsing in water. Chlorhydric water (0.5%) was then added for a few seconds to stain specimens in pink before another cycle of lithium carbonate. Fuschin Ponceau was then added for 5 min, and phosphomolybdic acid was added for 10 s before 5 min incubation. Lastly, a Light Green staining was done for 5 min, followed by acetified water for 30 s. Slides were finally dehydrated and mounted as depicted above. Final acquisition of images was done using the NanoZoomer Digital Pathology software (Hamamatsu, Massy, Japan). For immunofluorescence *per se*, the Discovery Ultra device (Roche, Illkirch, France) was used. Paraffin removal was done for 8 min (75°C) using the EZPrep reagent (Roche, Illkirch, France), followed by antigen retrieval (8 min / 95°C then 28 min / 100°C in Tris EDTA pH 8.0), followed by incubations of primary antibodies (**Suppl. Table 12)**, prior to washing three times with PBS / 0.1% Tween and incubation of secondary antibodies (60 min, 37°C, see figure legend) and identical washing. The Discovery DCC Kit 455 (Roche), the Discovery FAM Kit 505 (Roche), the Discovery Cy5 Kit 660 (Roche), and the Opal Polaris 780 Reagent (Akoya) were used. Counterstaining was performed using DAPI (250 ng/mL) before mounting on SuperFrost slides (Euromedex, Souffelweyersheim, France). Final acquisition of images was done using the NanoZoomer Digital Pathology software (Hamamatsu, Massy, France).

**Statistics**

All statistical analyses and statistical tests pertaining to the bioinformatics analysis were carried out with the R software (version 3.6.1). Heatmaps were generated with ComplexeHeatmap, principal component analysis was completed with ade4, and plotted with factoextra or ggplot. Gaussian finite models were performed with Cluster and ClusterProfiler. Figures were created using the R software. Statistics on samples derived from the French National HCC biobank were done as follows. Normal distribution of data was first assassed using the Shapiro-Wilk test. The associations between receptor transcript levels and clinico-pathological variables were determined in multiple comparison by Kruskal-Wallis test, Mann-Whitney test and Spearman correlation. We considered a Bonferroni-corrected *p*-value of 0.05/30 variables = 0.0017 as the threshold. Kaplan-Meier plots and log-rank tests were used to evaluate the prognostic value of the receptors by univariate Cox proportional hazards regression model.
