## Supplementary material for "Hepatocellular carcinoma hosts immature neurons and cholinergic tumors that correlate with adverse molecular features and outcomes": Suppl Table 1

**Supplementary Table 1. Characteristics of paired F4/HCC samples used in the study.** Median ± SD is shown for all qualitative data

| Variable | Available data (Total: 166) | Values |
| --- | --- | --- |
| Age, *y* ± SD | 166 | 69 ± 10 |
| Gender (male/female) | 166 | 140 (84%)/26 (16%) |
| Etiology (%) | 166 |  |
| Alcoholic liver disease |  | 36 (21%) |
| Hepatitis B virus |  | 46 (28%) |
| Hepatitis C virus |  | 39 (24%) |
| Non-alcoholic steato-hepatitis |  | 45 (27%) |
| Serum alpha-fetoprotein, >100 ng/mL | 111 | 17 (16%) |
| Child-Pugh score (A/B/C) (%) | 87 | 72 (83%)/9 (11%)/6 (6%) |
| Prothrombin, % ± SD | 146 | 81 ± 22 |
| Bilirubin, µMol/L ± SD | 137 | 14 ± 75 |
| Albumin, g/L ± SD | 100 | 36 ± 7 |
| Platelet count, G/L ± SD | 149 | 131 ± 102 |
| Encephalopathy (%) | 164 | 18 (11%) |
| Ascites (%) | 164 | 37 (3%) |
| Jaundice (%) | 162 | 22 (14%) |
| Esophageal varices (%) | 156 | 62 (40%) |
| Histological and gross features of the tumors |  |  |
| Tumor size, *mm* ± SD | 165 | 34 ± 31 |
| Tumor localization (left liver/right liver/double) | 165 | 47 (29%)/112 (68%)/6 (4%) |
| Intact tumor capsule (%) | 143 | 91 (64%) |
| Satellite nodules (%) | 165 | 34 (21%) |
| Macrovascular invasion, Microvascular invasion (%) | 151 | 22 (15%), 81 (52%) |
| Differentiation grade (poor/moderately/well) (%) | 165 | 15 (9%)/74 (45%)/76 (46%) |
| Architectural pattern (%) | 136 |  |
| Trabecular |  | 104 (76%) |
| Pseudoglandular |  | 10 (7%) |
| Compact |  | 5 (4%) |
| Clear cells |  | 4 (3%) |
| Others |  | 13 (9%) |
| Tumoral steatosis (%) | 165 | 108 (66%) |
| Tumoral necrosis (%) | 150 | 84 (56%) |
| Liver fibrosis score METAVIR (F4) (%) | 166 | 166 (100%) |
| Inflammation activity score (%) | 137 |  |
| METAVIR 0 |  | 51 (37%) |
| METAVIR 1 |  | 55 (40%) |
| METAVIR 2 |  | 23 (17%) |
| METAVIR 3 |  | 8 (6%) |
