## Supplementary material for "Hepatocellular carcinoma hosts immature neurons and cholinergic tumors that correlate with adverse molecular features and outcomes": Suppl Table 2

**Supplementary Table 2.**

**Significance assessment of neural features in HCC.** Significance calculated after Western blot and subsequent quantification (stratification based on etiology or Edmonson differentiation grade). Mann-Whitney test (n = 51 patients total, * p < 0.05, ** p < 0.01, *** p < 0.001).

| Comparison *tumoral* over *peritumoral* | HBV | HCV | NASH | ALD |
| --- | --- | --- | --- | --- |
| NeuN | ns | ns | ns | ns |
| DCX | * | ns | ns | ns |
| INA | ns | ns | ns | ns |
| TH | ns | ns | ** | ns |
| VAChT | ns | ns | ns | ns |
| VAChT/TH ratio | ns | ns | ns | ns |

| Comparison *high* over *low-medium* | HBV | HCV | NASH | ALD |
| --- | --- | --- | --- | --- |
| NeuN | * | ns | ns | ns |
| DCX | ns | ns | ns | ns |
| INA | ns | ns | * | * |
| TH | ns | ns | ns | ns |
| VAChT | ns | ns | * | ns |
| VAChT/TH ratio | ns | ns | ns | ns |
