## Supplementary material for "Hepatocellular carcinoma hosts immature neurons and cholinergic tumors that correlate with adverse molecular features and outcomes": Suppl Table 9

**Supplementary Table 9**

| **Adrenergic tumor pathways** | | **Cholinergic tumor pathways** | |
| --- | --- | --- | --- |
| **Prognosis agreement over adrenergic transcriptomic features** | **Prognosis discordance with over adrenergic transcriptomic**  **features** | **Prognosis agreement over cholinergic transcriptomic features** | **Prognosis discordance over cholinergic transcriptomic features** |
| CHIANG_LIVER_CANCER_SUBCLASS_PROLIFERATION_DN | CHIANG_LIVER_CANCER_SUBCLASS_INTERFERON_DN | CHIANG_LIVER_CANCER_SUBCLASS_PROLIFERATION_UP | None |
| Padj = 8.2E-117 | Padj = 8.3E-4 | Padj = 2.2E-90 |  |
| HOSHIDA_LIVER_CANCER_SUBCLASS_S3 |  | DESERT_STEM_CELL_HEPATOCELLULAR_CARCINOMA_SUBCLASS_UP |  |
| Padj = 1.5E-75 |  | Padj = 9.7E-35 |  |
| LEE_LIVER_CANCER_SURVIVAL_UP |  | VILLANUEVA_LIVER_CANCER_KRT19_UP |  |
| Padj = 1.4E-53 |  | Padj = 1.6E-26 |  |
| CHIANG_LIVER_CANCER_SUBCLASS_CTNNB1_UP |  | WOO_LIVER_CANCER_RECURRENCE_UP |  |
| Padj = 5.7E-49 |  | Padj = 2.2E-19 |  |
| WOO_LIVER CANCER RECURRENCE_DN |  | LEE_LIVER_CANCER_SURVIVAL_DN |  |
| Padj = 5.58E-42 |  | Padj = 5.3E-8 |  |
| VILLANUEVA_LIVER_CANCER_KRT19_DN |  | CHIANG_LIVER_CANCER_SUBCLASS_CTNNB1_DN |  |
| Padj = 1.1E-32 |  | Padj = 1.3E-6 |  |
| DESERT_PERIPORTAL_HEPATOCELLULAR_CARCINOMA_SUBCLASS_UP |  | HOSHIDA_LIVER_CANCER_SUBCLASS_S1 |  |
| Padj = 3.9E-28 |  | Padj = 7.3E-5 |  |
| BOYAULT_LIVER_CANCER_SUBCLASS_G123_DN |  | HOSHIDA_LIVER_CANCER_SURVIVAL_UP |  |
| Padj = 9.2E-27 |  | Padj = 8.7E-3 |  |
| ANDERSEN_LIVER_CANCER_KRT19_DN |  |  |  |
| Padj = 6.5E-24 |  |  |  |
| YAMASHITA_LIVER_CANCER_STEM_CELL_DN |  |  |  |
| Padj = 2.6E-24 |  |  |  |
| KIM_LIVER_CANCER_POOR_SURVIVAL_DN |  |  |  |
| Padj = 2.3E-21 |  |  |  |
| BOYAULT_LIVER_CANCER_SUBCLASS_G6_UP |  |  |  |
| Padj = 4.0E-16 |  |  |  |
| BOYAULT_LIVER_CANCER_SUBCLASS_G1_DN |  |  |  |
| Padj = 5.8E-9 |  |  |  |
| CHIANG_LIVER_CANCER_SUBCLASS_POLYSOMY7_UP |  |  |  |
| Padj = 8.3E-9 |  |  |  |
| HOSHIDA_LIVER_CANCER_SURVIVAL_DN |  |  |  |
| Padj = 3.3E-8 |  |  |  |
| BOYAULT_LIVER_CANCER_SUBCLASS_G12_DN |  |  |  |
| Padj = 2.0E-6 |  |  |  |
| HOSHIDA_LIVER_CANCER_LATE_RECURRENCE_DN |  |  |  |
| Padj = 9.0E-3 |  |  |  |
| BOYAULT_LIVER_CANCER_SUBCLASS_G56_UP |  |  |  |
| Padj = 1.8E-3 |  |  |  |
