## Supplementary material for "Hepatocellular carcinoma hosts immature neurons and cholinergic tumors that correlate with adverse molecular features and outcomes": Suppl Table 10

**Supplementary Table 10.**

**Biological descriptions of prognosis pathways associated with both HCC neuroclasses.**

| **Standard name** | **Brief description** | **Systematic name** |
| --- | --- | --- |
| ANDERSEN_LIVER_CANCER_KRT19_DN | Genes under-expressed in KRT19-positive [GeneID=3880] hepatocellular carcinoma. | M424 |
| BOYAULT_LIVER_CANCER_SUBCLASS_G1_DN | Down-regulated genes in hepatocellular carcinoma (HCC) subclass G1, defined by unsupervised clustering | M1883 |
| BOYAULT_LIVER_CANCER_SUBCLASS_G12_DN | Down-regulated genes in hepatocellular carcinoma (HCC) subclass G12, defined by unsupervised clustering | M12228 |
| BOYAULT_LIVER_CANCER_SUBCLASS_G123_DN | Down-regulated genes in hepatocellular carcinoma (HCC) subclass G123, defined by unsupervised clustering. | M2218 |
| BOYAULT_LIVER_CANCER_SUBCLASS_G6_UP | Up-regulated genes in hepatocellular carcinoma (HCC) subclass G6, defined by unsupervised clustering. | M4342 |
| CHIANG_LIVER_CANCER_SUBCLASS_CTNNB1_DN | [Top 200 marker genes down-regulated in the 'CTNNB1' subclass of hepatocellular carcinoma (HCC); characterized by activated CTNNB1 [GeneID=1499].](http://www.ncbi.nlm.nih.gov/gene/1499) | M8689 |
| CHIANG_LIVER_CANCER_SUBCLASS_CTNNB1_UP | [Top 200 marker genes up-regulated in the 'CTNNB1' subclass of hepatocellular carcinoma (HCC); characterized by activated CTNNB1 [GeneID=1499].](http://www.ncbi.nlm.nih.gov/gene/1499) | M16496 |
| CHIANG_LIVER_CANCER_SUBCLASS_INTERFERON_DN | All marker genes down-regulated in the 'interferon' subclass of hepatocellular carcinoma (HCC). | M14353 |
| CHIANG_LIVER_CANCER_SUBCLASS_POLYSOMY7_UP | Marker genes up-regulated in the 'chromosome 7 polysomy' subclass of hepatocellular carcinoma (HCC); characterized by polysomy of chromosome 7 and by a lack of gains of chromosome 8q. | M834 |
| CHIANG_LIVER_CANCER_SUBCLASS_PROLIFERATION_DN | [Top 200 marker genes down-regulated in the 'proliferation' subclass of hepatocellular carcinoma (HCC); characterized by increased proliferation, high levels of serum AFP [GeneID=174], and chromosomal instability.](http://www.ncbi.nlm.nih.gov/gene/174) | M16932 |
| CHIANG_LIVER_CANCER_SUBCLASS_PROLIFERATION_UP | [Top 200 marker genes up-regulated in the 'proliferation' subclass of hepatocellular carcinoma (HCC); characterized by increased proliferation, high levels of serum AFP [GeneID=174], and chromosomal instability.](http://www.ncbi.nlm.nih.gov/gene/174) | M3268 |
| DESERT_PERIPORTAL_HEPATOCELLULAR_CARCINOMA_SUBCLASS_UP | Genes up-regulated in the periportal-type subclass of hepatocellular carcinomas. | M34031 |
| DESERT_STEM_CELL_HEPATOCELLULAR_CARCINOMA_SUBCLASS_UP | Genes up-regulated in the stem cell-type subclass of hepatocellular carcinomas. | M34034 |
| HOSHIDA_LIVER_CANCER_LATE_RECURRENCE_DN | Genes whose expression correlated with lower risk of late recurrence of hepatocellular carcinoma (HCC). | M13658 |
| HOSHIDA_LIVER_CANCER_SUBCLASS_S1 | Genes from 'subtype S1' signature of hepatocellular carcinoma (HCC): aberrant activation of the WNT signaling pathway. | M5311 |
| HOSHIDA_LIVER_CANCER_SUBCLASS_S3 | Genes from 'subtype S3' signature of hepatocellular carcinoma (HCC): hepatocyte differentiation. | M1286 |
| HOSHIDA_LIVER_CANCER_SURVIVAL_DN | Survival signature genes defined in adjacent liver tissue: genes correlated with good survival of hepatocellular carcinoma (HCC) patients. | M5451 |
| HOSHIDA_LIVER_CANCER_SURVIVAL_UP | Survival signature genes defined in adjacent liver tissue: genes correlated with poor survival of hepatocellular carcinoma (HCC) patients. | M6939 |
| KIM_LIVER_CANCER_POOR_SURVIVAL_DN | Genes under-expressed in hepatocellular carcinoma (HCC) with poor survival | M534 |
| LEE_LIVER_CANCER_SURVIVAL_DN | Genes highly expressed in hepatocellular carcinoma with worse survival. | M7987 |
| LEE_LIVER_CANCER_SURVIVAL_UP | Genes highly expressed in hepatocellular carcinoma with better survival. | M6145 |
| VILLANUEVA_LIVER_CANCER_KRT19_DN | Genes under-expressed in KRT19-positive [GeneID=3880] hepatocellular carcinoma (HCC). | M373 |
| VILLANUEVA_LIVER_CANCER_KRT19_UP | Genes over-expressed in KRT19-positive [GeneID=3880] hepatocellular carcinoma (HCC). | M336 |
| WOO_LIVER CANCER RECURRENCE_DN | Genes negatively correlated with recurrence free survival in patients with hepatitis B-related (HBV) hepatocellular carcinoma (HCC). | M9911 |
| WOO_LIVER_CANCER_RECURRENCE_UP | Genes positively correlated with recurrence free survival in patients with hepatitis B-related (HBV) hepatocellular carcinoma (HCC). | M12602 |
| YAMASHITA_LIVER_CANCER_STEM_CELL_DN | Genes down-regulated in hepatocellular carcinoma (HCC) cells with hepatic stem cell properties. | M9206 |
