## Supplementary material for "Hepatocellular carcinoma hosts immature neurons and cholinergic tumors that correlate with adverse molecular features and outcomes": Suppl Table 13

**Supplementary Table 13.**

Antibodies used in this study.

| **Antigen** | **Ig Species** | **Cat. Number & supplier** | **Antibody registry (RRID) #** |
| --- | --- | --- | --- |
| Beta-tubulin | Rabbit polyclonal | Ab6046, Abcam | AB_2210370 |
| NeuN | Mouse monoclonal | MAB377, Millipore | AB_2298772 |
| DCX | Mouse monoclonal  Rabbit polyclonal | Ab18723, Abcam (WB)  MABN707, Millipore (IHC) | AB_732011  ND |
| Alpha-internexin | Mouse monoclonal | MAB5224, Millipore | AB_2127486 |
| TH | Rabbit polyclonal  Rabbit polyclonal | AB152, Millipore (WB)  Ab112, Abcam (IHC) | AB_390204  AB_297840 |
| VAChT | Mouse monoclonal  Rabbit polyclonal | SAB5200240, Sigma (WB)  PA5-85782, Invitrogen (IHC) | ND  AB_2992918 |
| Anti-mouse-HRP | Goat polyclonal | A4416, Sigma | AB_258167 |
| Anti-Rabbit-HRP | Goat polyclonal | A6154, Sigma | AB_258284 |
