## Supplementary material for "Hepatocellular carcinoma hosts immature neurons and cholinergic tumors that correlate with adverse molecular features and outcomes": Suppl Table 14

**Supplementary Table 14.**

**Human qPCR primers.**

| **Gene symbol** | **Genbank**  **acc. #** | **Primer sequences (5’-3’, F/R)** | **PCR conditions** | **Amplicon**  **length, bp** |
| --- | --- | --- | --- | --- |
| *GUS* | NM_001293105 | CGTGGTTGGAGAGCTCATTTGGAA  TTCCCCAGCACTCTCGTCGGT | Denaturing, 95°C;  annealing, 55°C | 72 |
| *CHRM3* | [NM_000740](http://www.ncbi.nlm.nih.gov/entrez/query.fcgi?cmd=Search&db=Nucleotide&term=NM_000740) | TACCTGGAACAGGTGGAGC  GATCCCGGCATAGGACAGAG | Denaturing, 95°C;  annealing, 60°C | 73 |
| *CHRNA4* | NM_000744.7 | CTCCGAGCTCATCTGGCG  TCCCCGTCAGCATTGTTGTA | Denaturing, 95°C;  annealing, 60°C | 72 |
| *CHRNA7* | NM_000746.6 | GCTGGTCAAGAACTACAATCCC  CTCATCCACGTCCATGATCTG | Denaturing, 95°C;  annealing, 60°C | 106 |
| *ADRA1A* | NM_000680.4 | CCAAGACGGATGGCGTTTG  TGGACACTGTAATCCTGGCAG | Denaturing, 95°C;  annealing, 60°C | 75 |
| *ADRA1B* | NM_000679.4 | TGGGGCGGATCTTCTGTGA  GTGACCAGCGTGGGATACTG | Denaturing, 95°C;  annealing, 60°C | 136 |
| *ADRA1D* | NM_000678.4 | GCCGCTCGGCTCCTTG  GGCTGGAACAGGGGTAGATG | Denaturing, 95°C;  annealing, 60°C | 116 |
| *ADRB1* | NM_000684.3 | ATCGAGACCCTGTGTGTCATT  GTAGAAGGAGACTACGGACGAG | Denaturing, 95°C;  annealing, 60°C | 267 |
| *ADRB2* | NM_000024.6 | TGGTGTGGATTGTGTCAGGC  GGCTTGGTTCGTGAAGAAGTC | Denaturing, 95°C;  annealing, 60°C | 128 |
| *ADRB3* | NM_000025.3 | GACCAACGTGTTCGTGACTTC  GCACAGGGTTTCGATGCTG | Denaturing, 95°C;  annealing, 60°C | 175 |
