## Supplementary material for "Hepatocellular carcinoma hosts immature neurons and cholinergic tumors that correlate with adverse molecular features and outcomes": Suppl Figures

Suppl. Figure 1

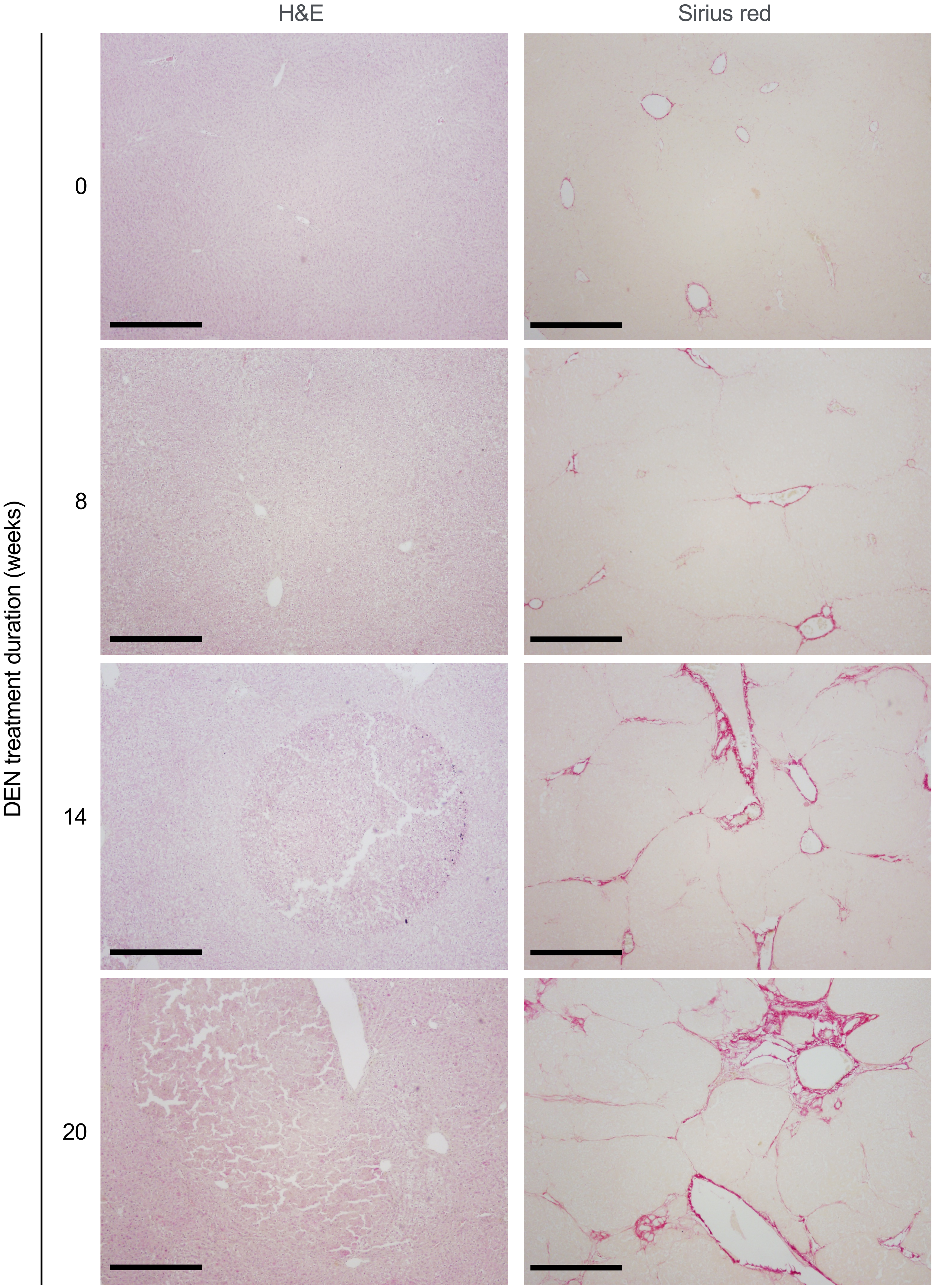

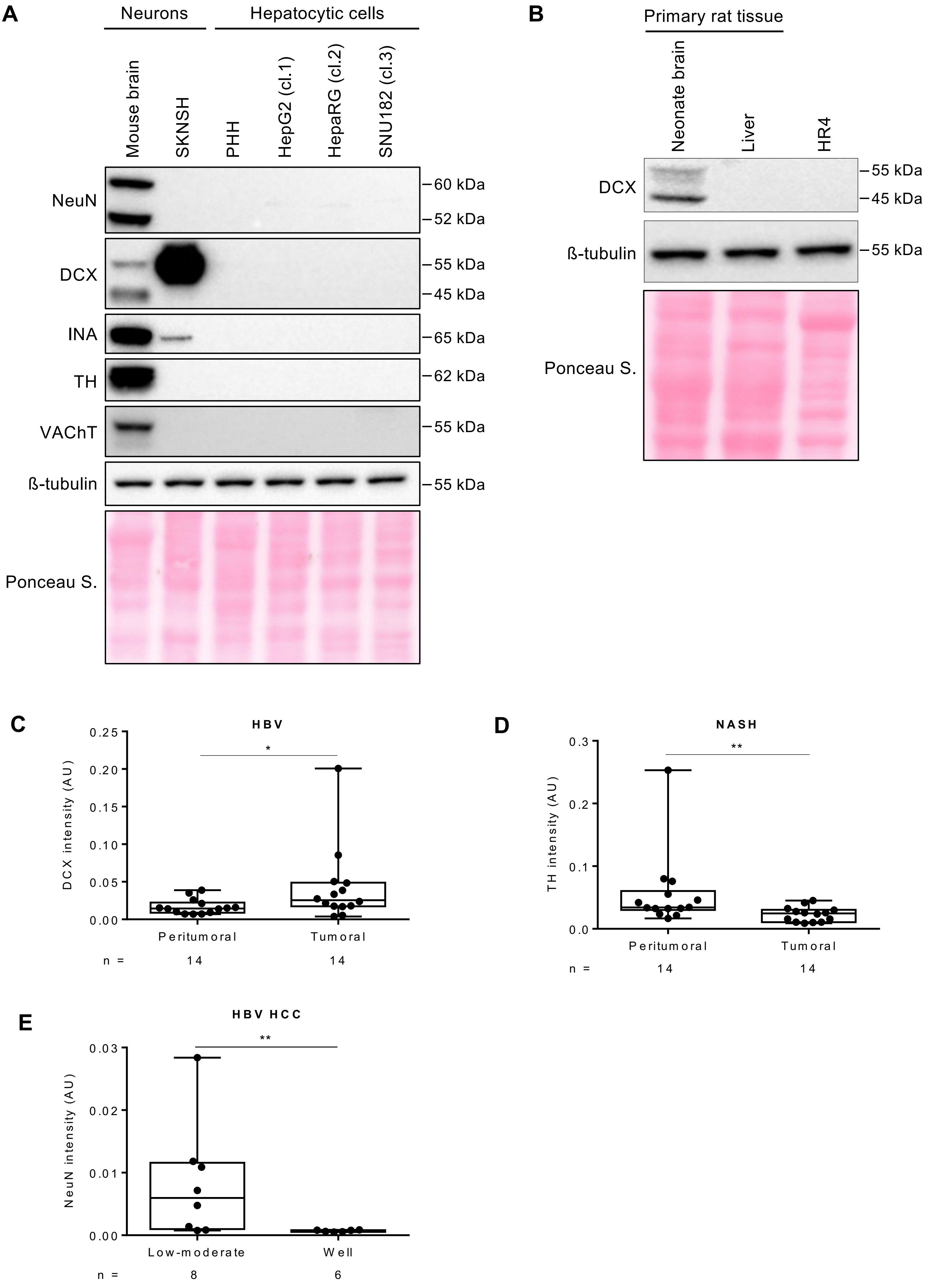
A
Neurons
Hepatocytic cells
Mouse brain
SKNSH
PHH
HepG2 (cl.1)
HepaRG (cl.2)
SNU182 (cl.3)
NeuN
DCX
INA
TH
VAChT
ß-tubulin
Ponceau S.
60 kDa
52 kDa
55 kDa
45 kDa
65 kDa
62 kDa
55 kDa
55 kDa
B
Primary rat tissue
Neonate brain
Liver
HR4
DCX
ß-tubulin
Ponceau S.
55 kDa
45 kDa
55 kDa
C
HBV
DCX intensity (AU)
0.25
0.20
0.15
0.10
0.05
0.00
*
Peritumoral
Tumoral
n =
14
14
D
NASH
TH intensity (AU)
0.3
0.2
0.1
0.0
**
Peritumoral
Tumoral
n =
14
14
E
HBV HCC
NeuN intensity (AU)
0.03
0.02
0.01
0.00
**
Low-moderate
Well
n =
8
6

Suppl. Figure 3

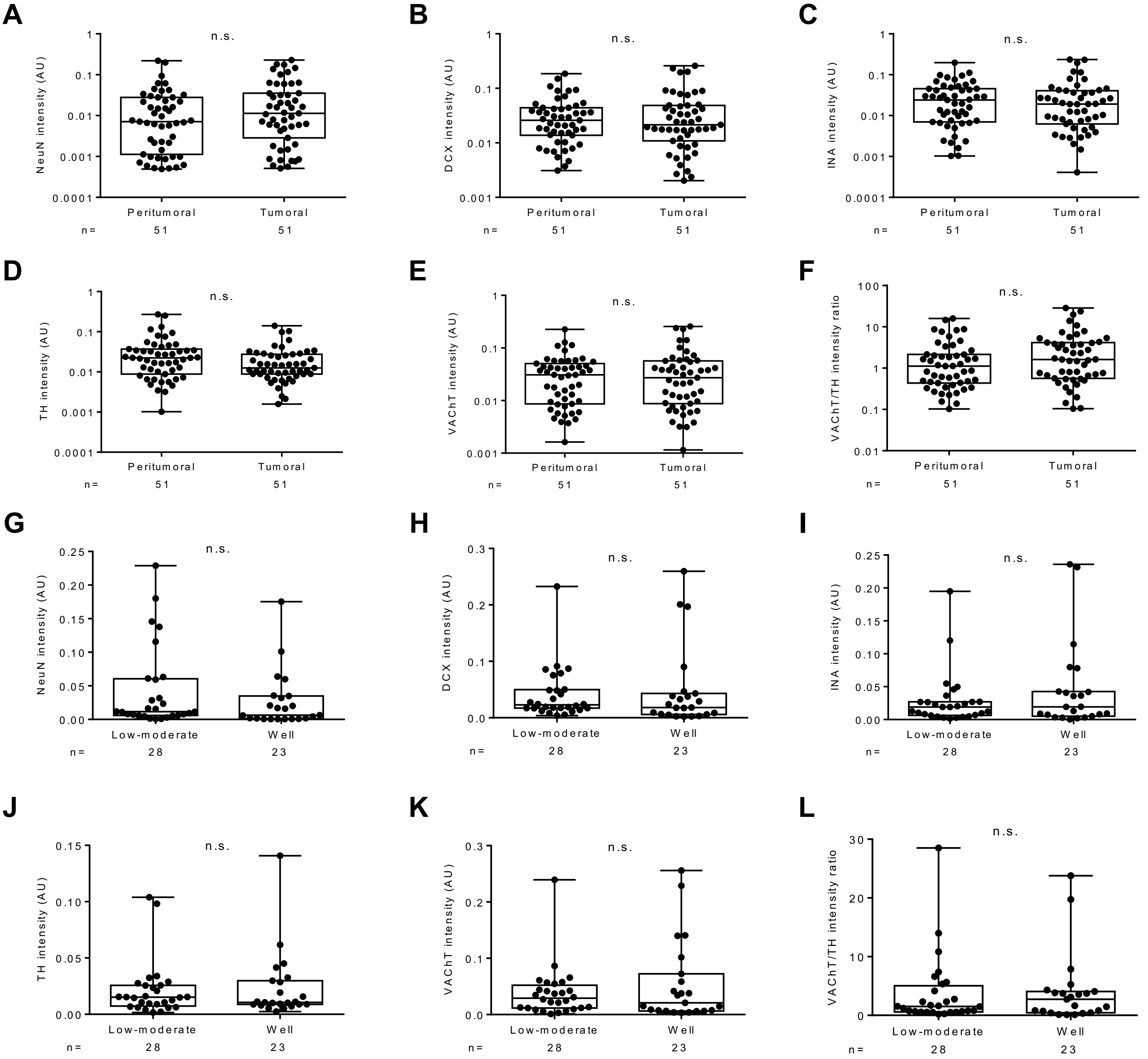

Suppl. Figure 4

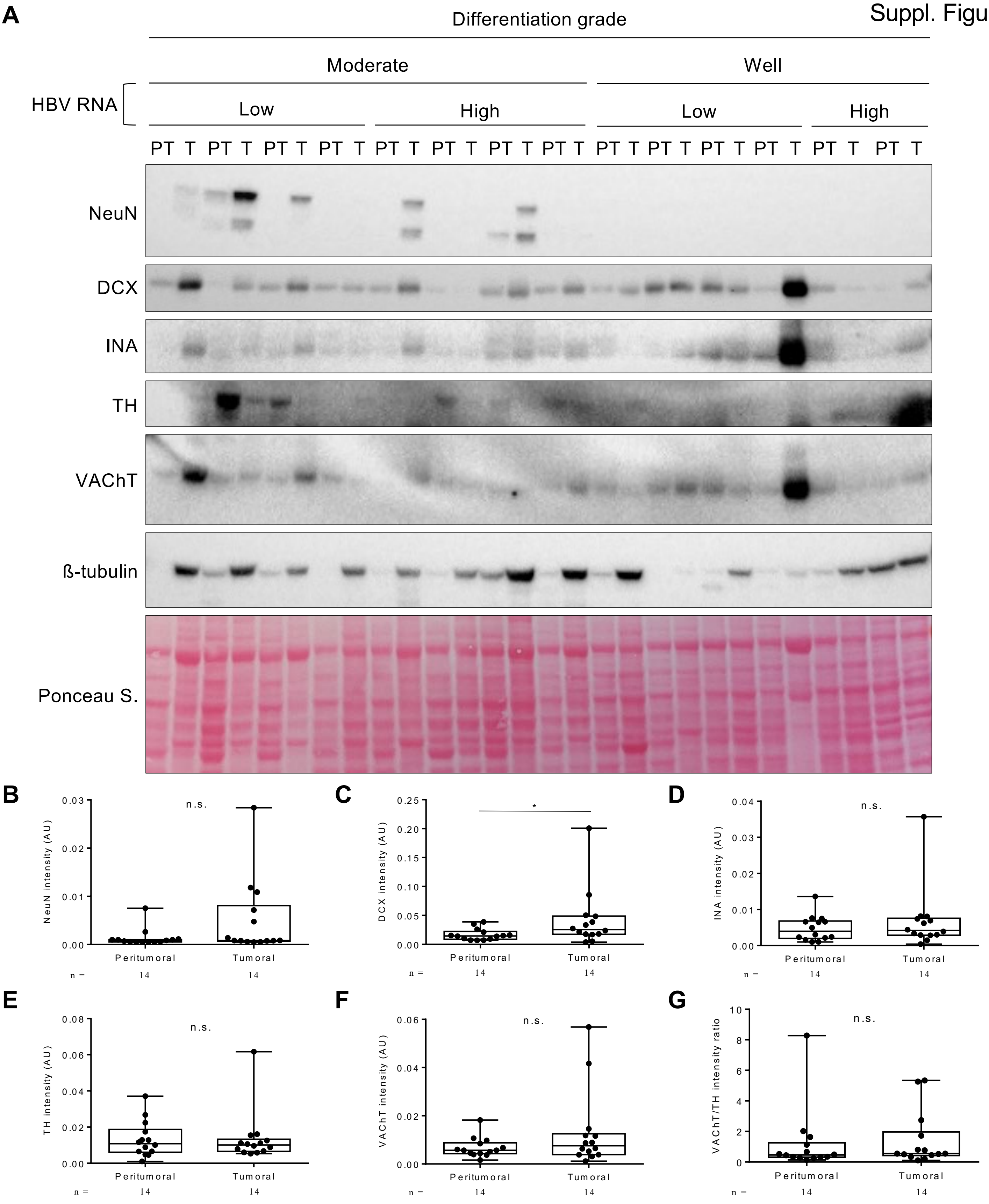

Suppl. Figure 5

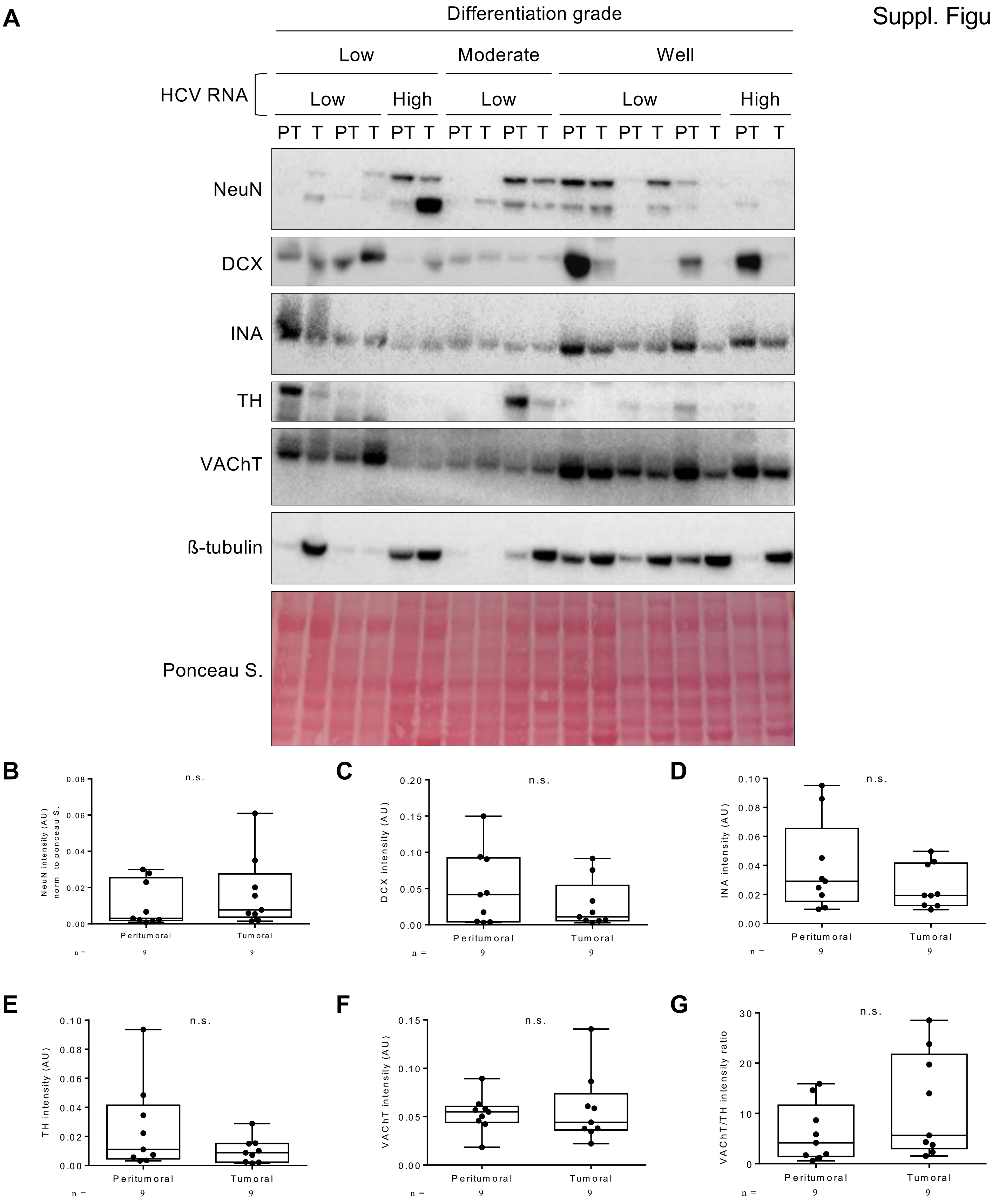

Suppl. Figure 6

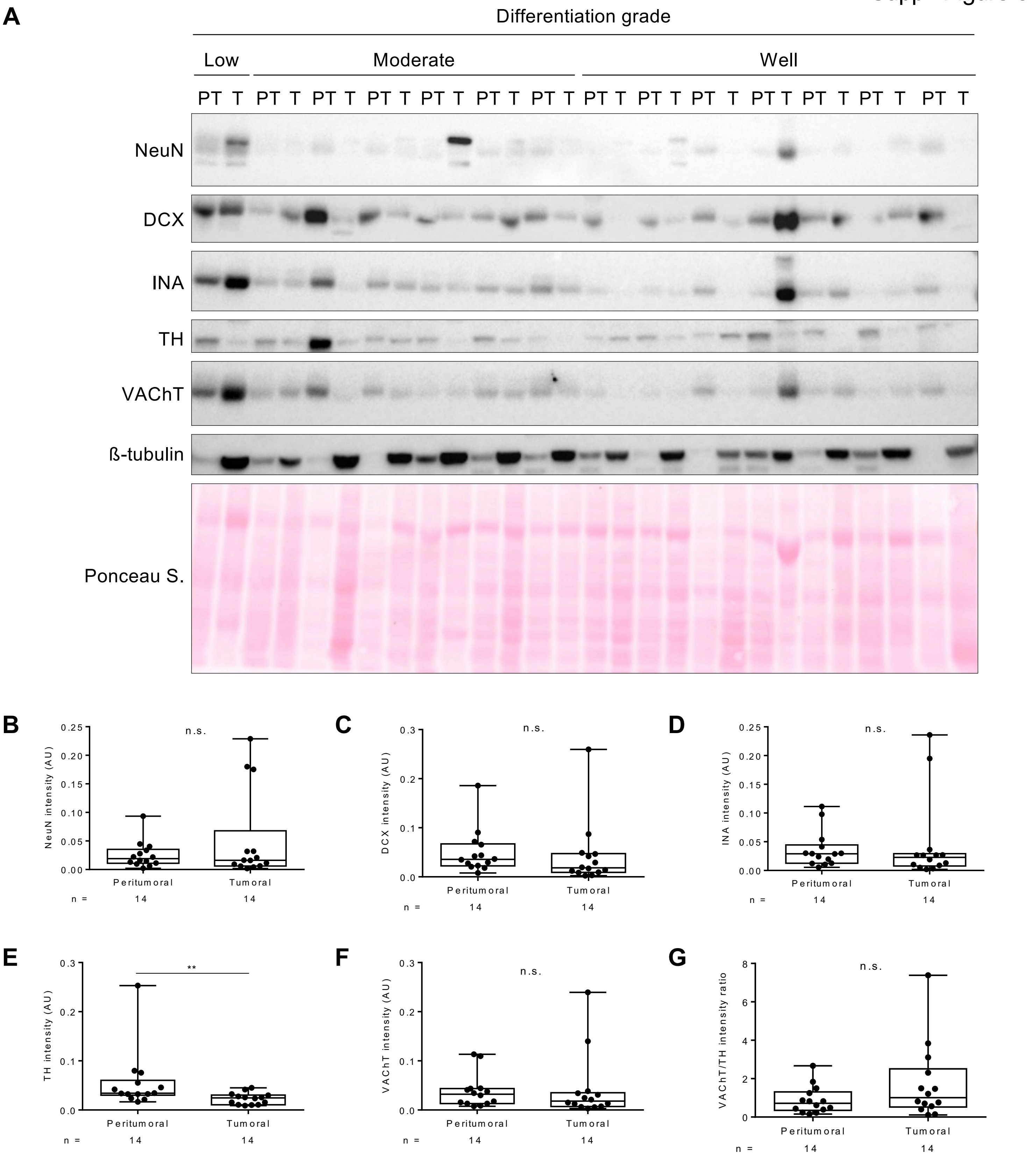

Suppl. Figure 7

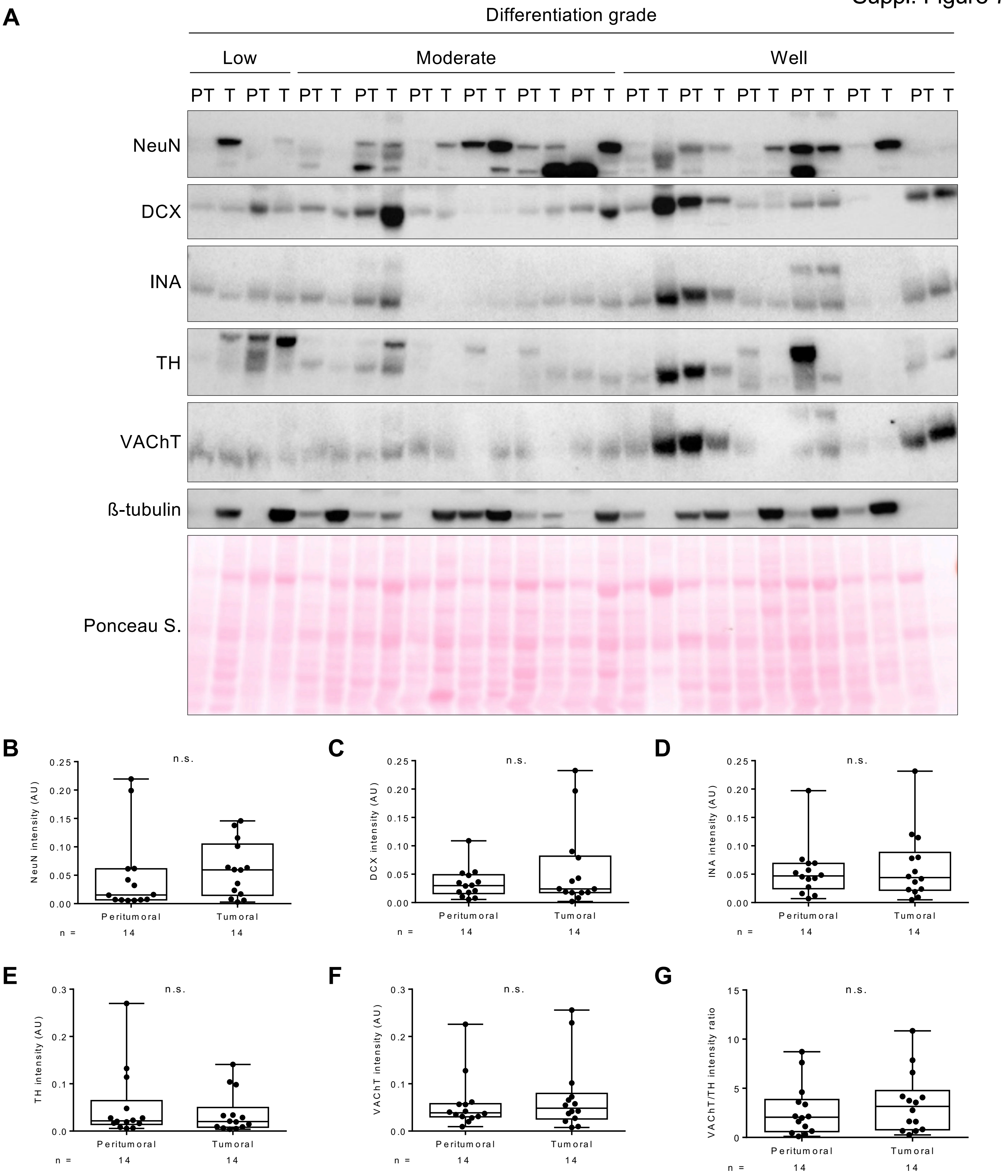

Suppl. Figure 8

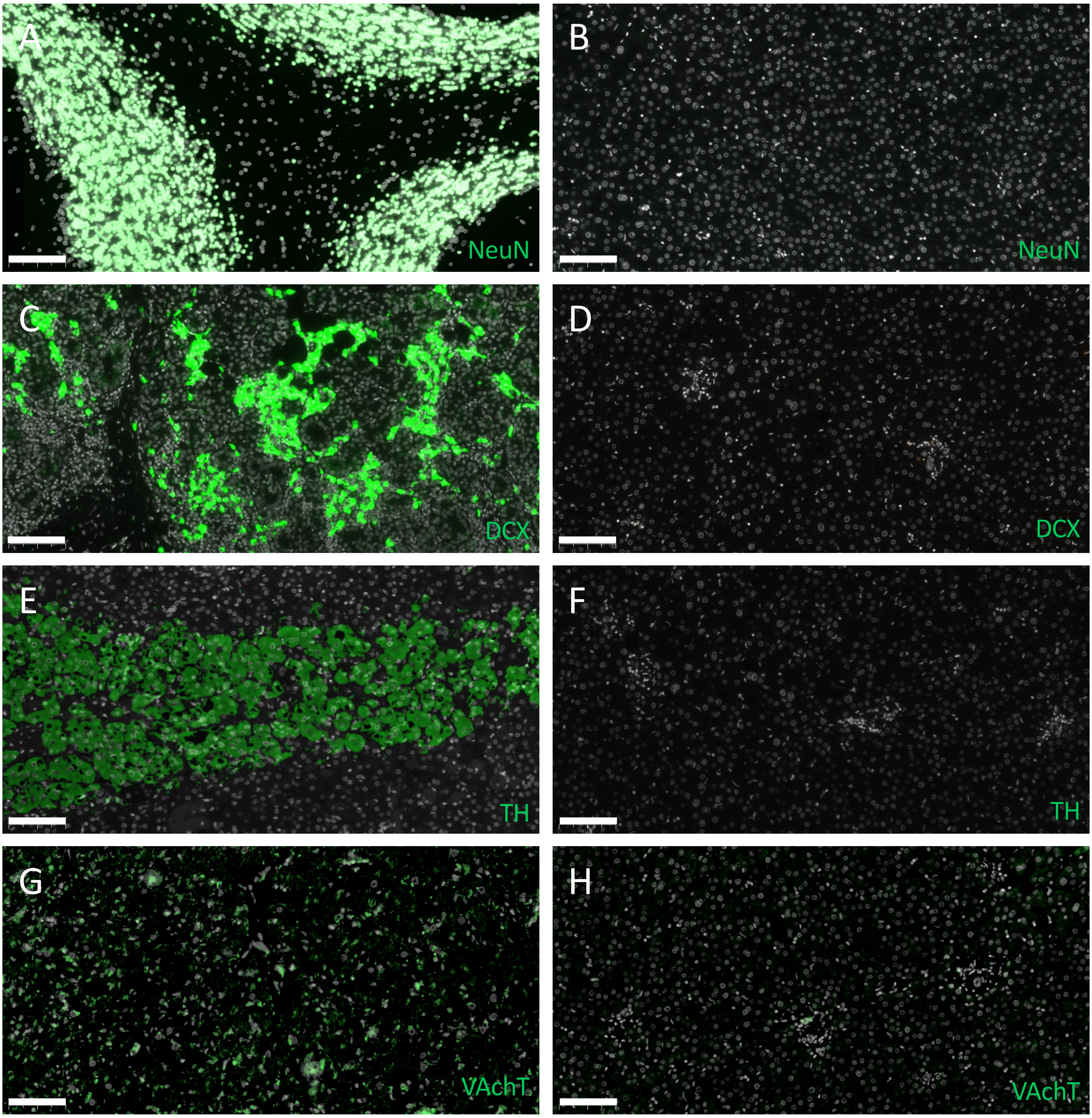

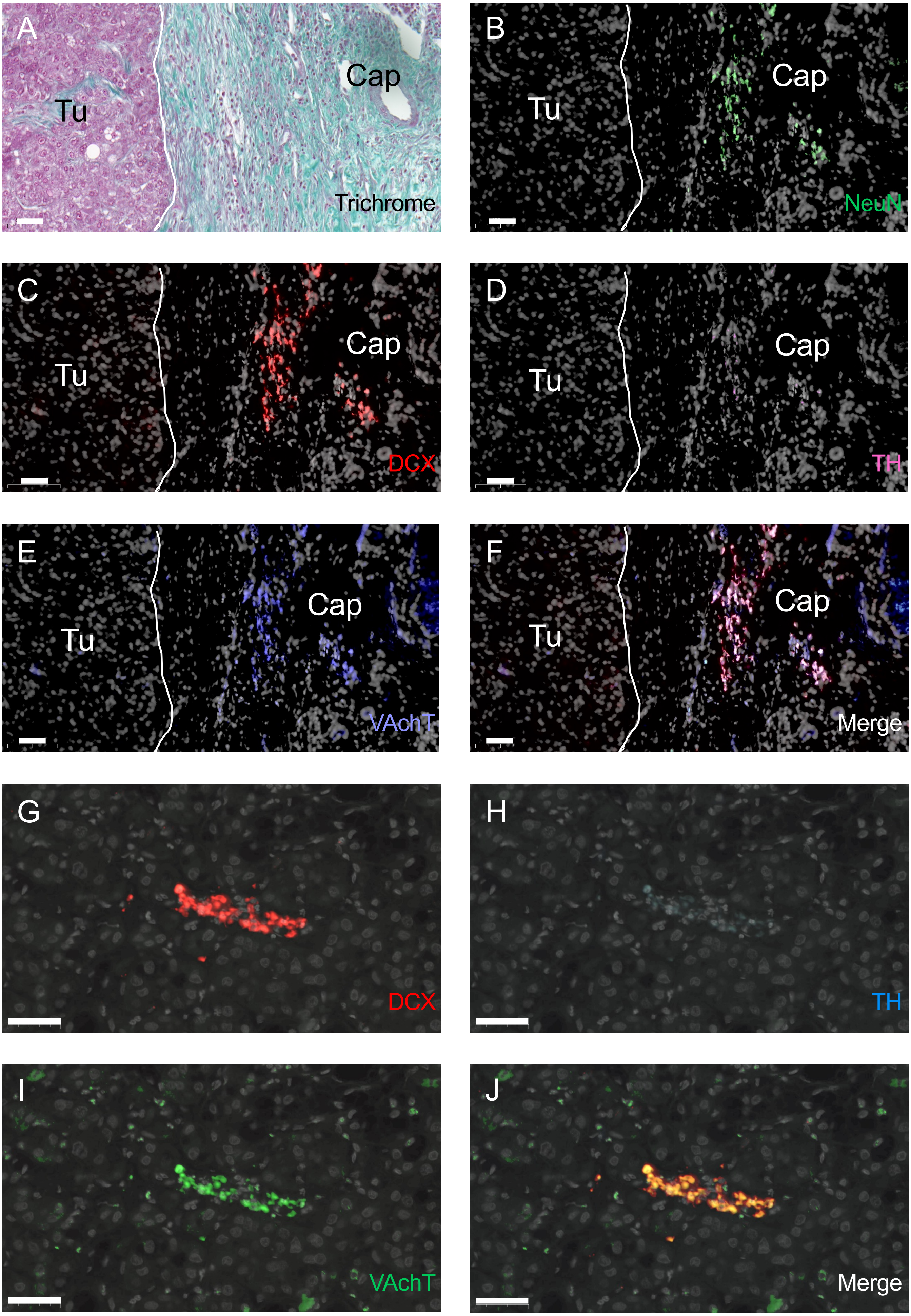
A
Tu
Cap
Trichrome
B
Tu
Cap
NeuN
C
Tu
Cap
DCX
D
Tu
Cap
TH
E
Tu
Cap
VAchT
F
Tu
Cap
Merge
G
DCX
H
TH
I
VAchT
J
Merge

Suppl. Figure 10

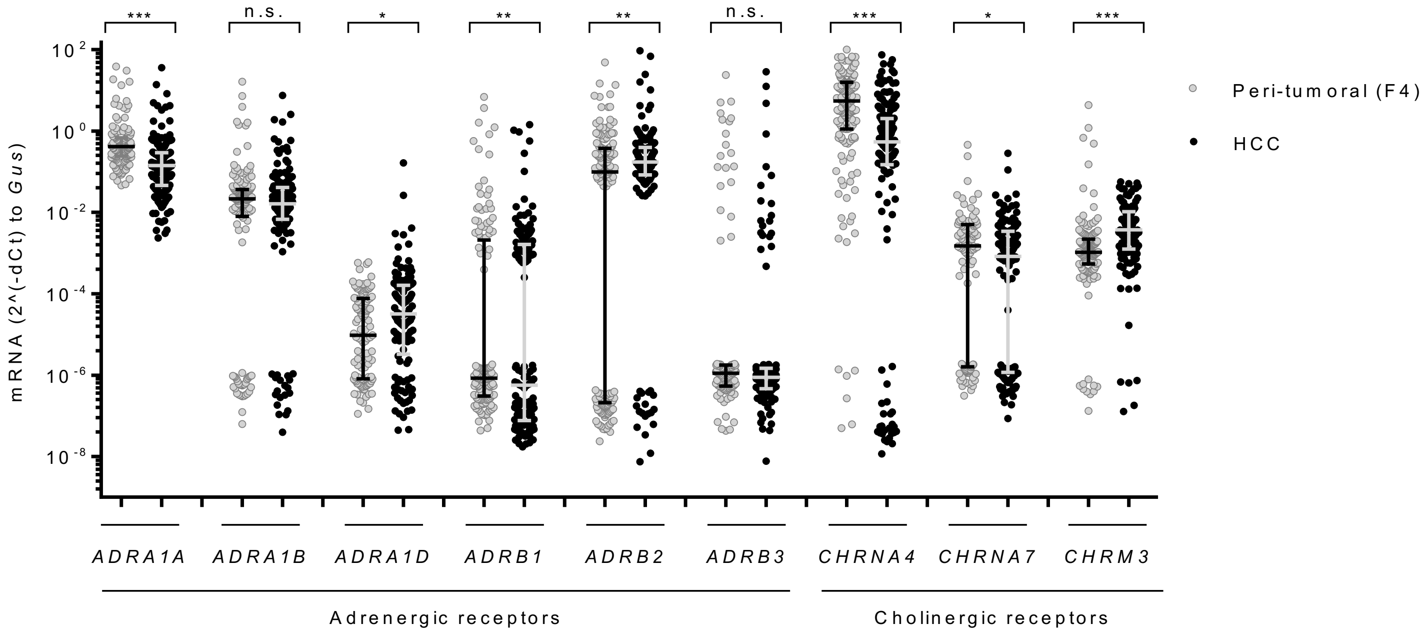

Suppl. Figure 11

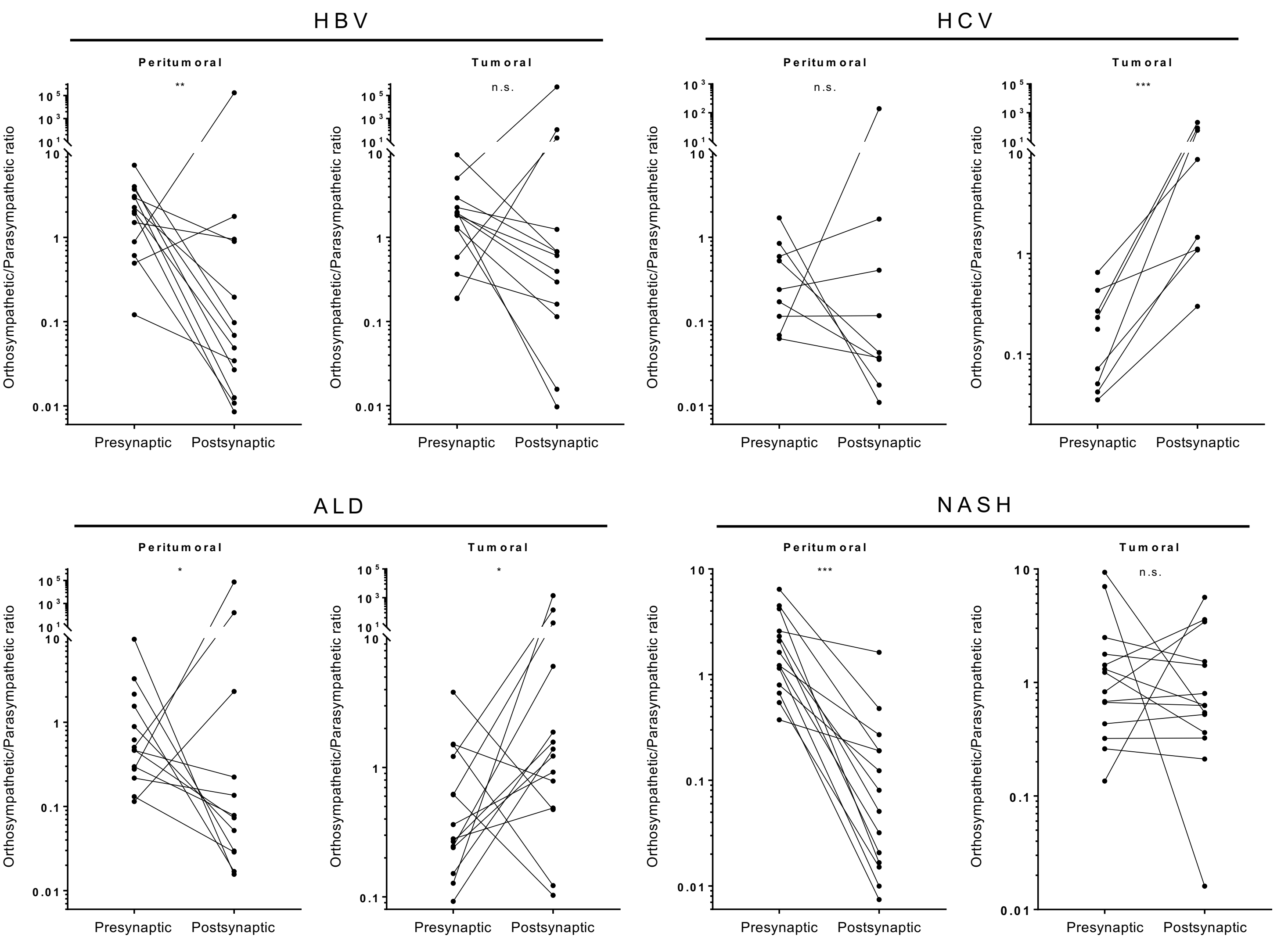

**A**

Suppl. Figure 12

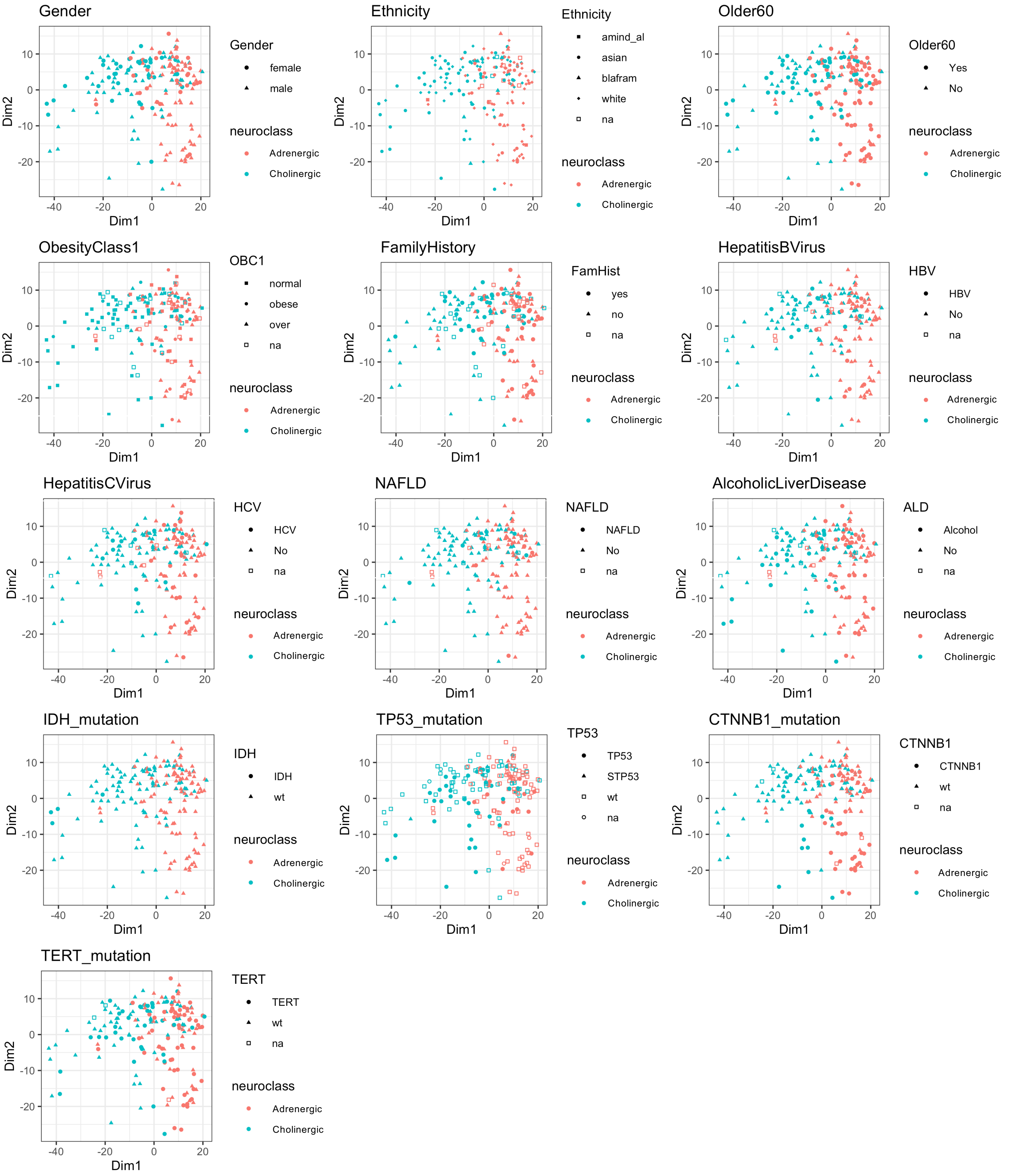
